# Translation dynamics in human cells visualized at high-resolution reveal cancer drug action

**DOI:** 10.1101/2023.03.02.529652

**Authors:** Huaipeng Xing, Reiya Taniguchi, Iskander Khusainov, Jan Philipp Kreysing, Sonja Welsch, Beata Turoňová, Martin Beck

**Affiliations:** Department of Molecular Sociology, Max Planck Institute of Biophysics, 60438 Frankfurt am Main, Germany; Faculty of Biochemistry, Chemistry and Pharmacy, Goethe University Frankfurt am Main, 60438 Frankfurt am Main, Germany; IMPRS on Cellular Biophysics, 60438 Frankfurt am Main, Germany; Central Electron Microscopy Facility, Max Planck Institute of Biophysics, 60438 Frankfurt am Main, Germany

## Abstract

Ribosomes catalyze protein synthesis by cycling through various functional states. These states have been extensively characterized in vitro, yet their distribution in actively translating human cells remains elusive. Here, we optimized a cryo-electron tomography-based approach and resolved ribosome structures inside human cells with a local resolution of up to 2.5 angstroms. These structures revealed the distribution of functional states of the elongation cycle, a Z tRNA binding site and the dynamics of ribosome expansion segments. In addition, we visualized structures of Homoharringtonine, a drug for chronic myeloid leukemia treatment, within the active site of the ribosome and found that its binding reshaped the landscape of translation. Overall, our work demonstrates that structural dynamics and drug effects can be assessed at near-atomic detail within human cells.

**One-Sentence Summary:** Snapshots of ribosome dynamics at near-atomic resolution within native and drug-treated human cells are revealed.

## Main Text

The eukaryotic ribosome (80S) consists of two subunits (60S and 40S) that translate messenger RNA (mRNA) into proteins (1). Purified 80S ribosomes were extensively studied under different conditions in vitro, which provided the molecular details of translation (2, 3). The translation process can be divided into four main stages: initiation, elongation, termination and ribosome recycling (3). During elongation, the rotation of the 40S, the association of elongation factors and the translocation of transfer RNAs (tRNAs) are coordinated to synthesize nascent chains (3). tRNAs have three canonical binding sites on the ribosome: aminoacyl (A), peptidyl (P), and exit (E) sites (3). A noncanonical tRNA binding site called the Z site, located in an extreme position past the E site, has been identified in structures of isolated translationally inactive mammalian ribosomes (4). It has been speculated that the Z site may represent a late-stage intermediate of tRNA ejection downstream of the E site with similarity to internal ribosome entry site (IRES) interactions, but the exact physiological relevance remains obscure because the possibility of isolation artifacts could not entirely be ruled out (4).

The 70S ribosome from *Mycoplasma pneumoniae* and 80S ribosome from *Dictyostelium discoideum* and the respective translation elongation processes were visualized inside cells using cryo-electron tomography (cryo-ET), revealing differences in translation elongation (*5*–*7*). For example, the most abundant elongation intermediate in bacterial *M. pneumoniae* cells was the ‘A, P’ state, in which tRNAs were identified in the respective positions. In contrast, the elongation factor bound ‘eEF1A, A/T, P’ state was most abundant in eukaryotic *D. discoideum* cells. In microsomes isolated from human cells, the membrane-associated elongation cycle was resolved, revealing the ‘eEF1A, A/T, P, E’ state as the most prominent (8). However, to which extent these insights are applicable to actively translating human cells remains elusive and is addressed in this study.

Homoharringtonine (HHT) is a natural compound that binds to the ribosome and inhibits protein synthesis (9). It is used to treat patients with chronic myeloid leukemia clinically and as a reference to study new anti-cancer ribosome inhibitors (9, 10). The HHT-bound ribosome structure was determined by incubating the purified 80S ribosome with HHT in vitro (11, 12), revealing the binding site at the peptidyl transferase center (PTC). In the cellular context, it remains unclear how HHT affects translation dynamics and how exactly it inhibits protein synthesis. In this study, we applied cryo-focused ion beam (cryo-FIB), cryo-ET and advanced data processing algorithms to determine the near-atomic structures of ribosomes and analyzed the abundance and organization of different ribosome states in untreated and HHT-treated human cells. We show that prominent intermediates of the human translation elongation cycle are distinct from the previously in vitro studies. They contain unique structural features and are fundamentally shifted by HHT.

### Ribosome is bound with HHT inside human cells

To study the 80S ribosome structures inside human cells, we first prepared cryo-FIB-milled lamellae from 35 native (untreated) human embryonic kidney (HEK) 293 cells and acquired 358 tilt-series at a pixel size of 1.223 Å at the specimen level (fig. S1A and table S1). We used template matching to identify ribosomes within the reconstructed tomograms (Fig. 1A, fig. S1B and movie S1). After classification and refinement, we determined the ribosome structure at an overall resolution of ∼3.2 Å and locally resolved features up to 2.5 Å in resolution from untreated cells (Fig. 1B and figs. S1 to S3). The density of ribosomal protein side chains, ribosomal RNA (rRNA) bases, tRNA bases and ions were well resolved (fig. S1E), indicating the good quality of the map. Using the same approach, we processed 352 tilt series from 32 HHT-treated cells. In the resulting structure, the density of HHT was visible (Fig. 1C and fig. S1, F to H) and the position of HHT at the PTC showed a high similarity to previous *in vitro* analysis (9, 11). The P-tRNA was poorly resolved compared to the untreated ribosome (Fig. 1C), suggesting that the translation was altered after HHT treatment.

**Fig. 1.**
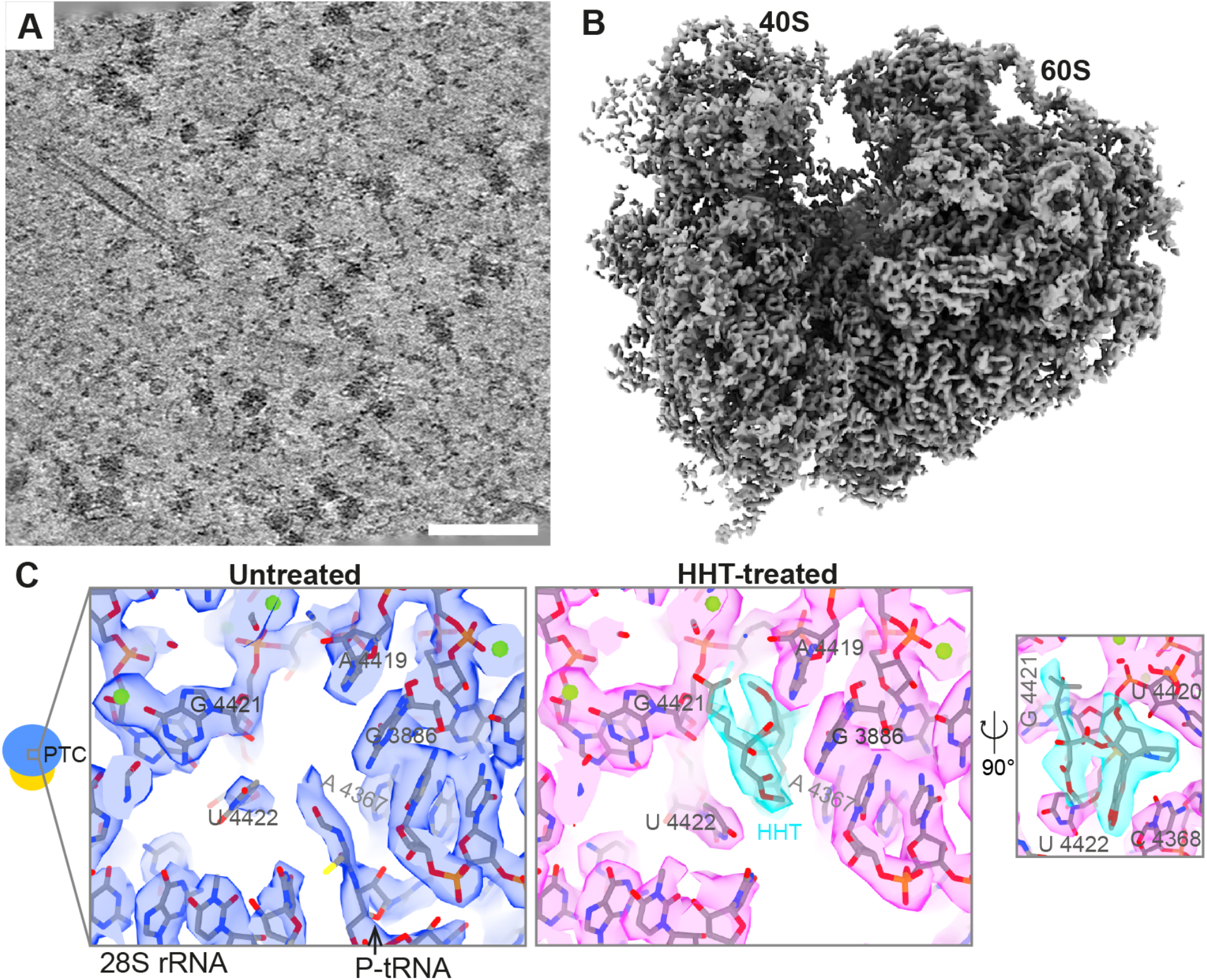
80S ribosome structures in human cells. (**A**) A tomographic slice from an untreated human cell. Scale bar, 100 nm. (**B**) Map of 80S ribosome from untreated cells. (**C**) The PTC of ribosomes from untreated and HHT-treated cells (see also movie S2). Black arrow, P-tRNA with a nascent chain. Green sphere, potential Mg^2+^.

### Translation elongation landscape in human cells

We next investigated the translation elongation process. In untreated cells, focused classification was performed with dedicated tRNA and elongation factor masks, resulting in eight ribosome states resolved from 3.4 Å to 16.4 Å (figs. S2 and S3 and table S2). These states were well-explained by previously reported structural models that captured various elongation states (fig. S4) (*4, 13*–*15*). Six classes, accounting for 85.5% of all identified ribosomes, were assigned to the translation elongation cycle based on their similarity to the previously reported structures (Fig. 2A) (3, 5). The elongation cycle is thought to start from the non-rotated ‘P’ state that was detected but not particularly abundant in our data (5) (Fig. 2A). The ‘eEF1A, A/T, P’ was the most prominent, consistent with *in situ* analysis of the lower eukaryote *Dictyostelium discoideum* (7). The global resolution of this state reached ∼3.4 Å and the local resolution was relatively uniform, suggesting its structural homogeneity (fig. S5A). The following states were non-rotated ‘A/T, P’ and ‘A, P’. The latter was less abundant as compared to bacteria, where it was the most populated (5), indicating the divergence across organisms. The subsequent state was identified as ‘eEF2, ap/P, E’ with rotated 40S, suggesting that ap-tRNA had not fully reached the ‘P’ site when the P-tRNA completely moved to the E site. Succeeding, there may be two possible sequences of events: either eEF2 and E-tRNA dissociate from the ribosome at first to form the ‘P’ state that subsequently binds to the eEF1A-tRNA, or the eEF1A-tRNA may displace eEF2 with subsequent departure of the E-tRNA (5) (Fig. 2A). The two decoding nucleotides A1824 and A1825 are thought to stabilize the A-site tRNA by flipped out of helix 44 of 18S rRNA and hydrogen bonding (PDB 5LZS) (16). As expected, these nucleotides were flipped out of the helix at the ‘A, P’ state (fig. S4K). This is however contrasted by the ‘eEF1a, A/T, T’ state (fig. S4K), indicating that the respective ribosomes may still be scanning for the correct tRNA inside the native cytoplasm of cells.

**Fig. 2.**
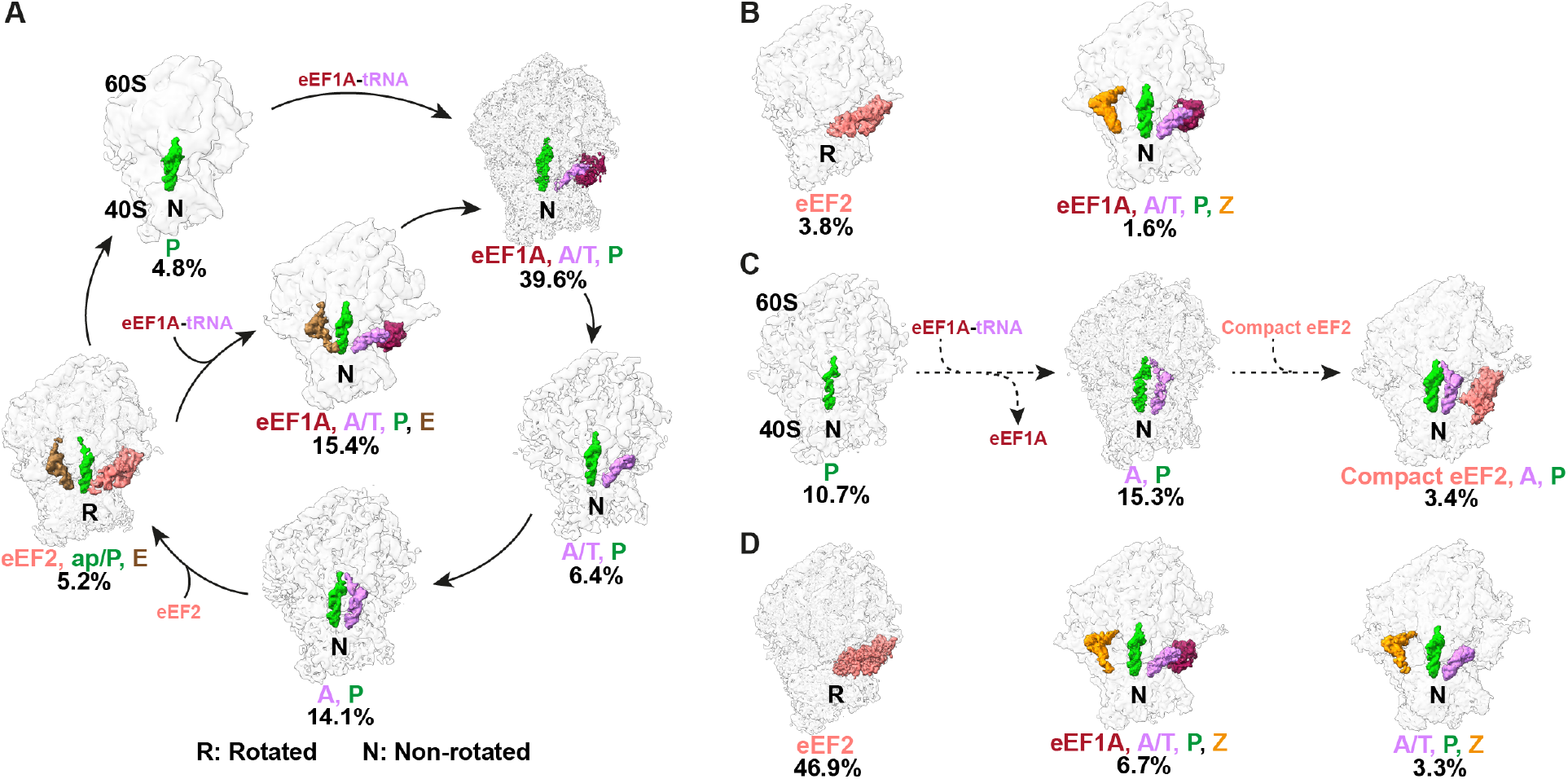
Ribosome states in native untreated and HHT-treated cells. (**A**) Six ribosome states are assigned to the elongation cycle in untreated cells. (**B**) The ‘eEF2’ and ‘eEF1A, A/T, P, Z’ states in untreated cells. (**C**) Three potential translation elongation intermediate states in the HHT-treated cells. The dashed arrow illustrates how the elongation states may connect. (**D**) The ‘eEF2’, ‘eEF1A, A/T, P, Z’ and ‘A/T, P, Z’ states inside the treated cells.

Two further states were identified that are non-obvious to place into the elongation cycle (Fig. 2B). The rotated ‘eEF2’ state without tRNA fitted the previous structural model very well (15, 17) (fig. S4G), although the eEF2 position was shifted by ∼1.7 nm as compared to the ‘eEF2, ap/P, E’ state (fig. S4I). The remaining class, which accounts for only 1.6% of all identified ribosomes (Fig. 2B), was consistent with the non-rotated ‘eEF1A, A/T, P, Z’ state that had been suggested to be rather transient (4). Although the resolution of this average was only moderate in untreated cells, the observed electron optical density at the Z site fitted well with the purified ribosome at the non-rotated ‘P, Z’ state (Fig. 2B and fig. S4, H and J) (4). To our knowledge, alternative factors binding to this site have not been identified. More importantly, we obtained higher resolution in HHT-treated cells (see below), which further strengthened this conclusion. Collectively, our data represented the detailed analysis of the translation landscape inside native human cells, which differs from previous *ex vivo* studies (8, 18), indicating the ribosome isolation process may affect our ability to capture native elongation intermediates (5, 7).

### HHT modifies the elongation landscape in an unanticipated manner

The anti-cancer drug HHT blocks the peptide-bond formation (9, 19). Previous in vitro analysis of purified ribosomes revealed that HHT binds in the A-site cleft in the PTC (11, 20). Upon HHT treatment, one would thus expect to observe a strong enrichment of ribosomes with the A and P-site tRNAs bound (11). Indeed, *M. pneumoniae* ribosomes treated with 0.5 mg/ml (1,547.4 µM) chloramphenicol (Cm), a related antibiotic binding to the PTC of the 50S, showed over 70% of ribosomes at the ‘A, P’ state (5, 21). The effect of HHT on human ribosome states in cells remains unknown. To address this, we exposed cells to HHT concentrations of 0.055 mg/ml (100 µM) in the medium, which resulted in a significantly lower cellular protein concentration than untreated cells (fig. S6A), implying that protein synthesis was inhibited. At the time point examined by cryo-EM, the ATP level, which serves as an indicator of cell viability, was not yet reduced (fig. S6B), and the cell morphology of untreated and treated cells was indistinguishable (fig. S6C). Classification, as described above, identified six states resolved into the 3.7 Å to 11.5 Å range (figs. S7 to S10 and table S3). Unexpectedly, the rotated state with eEF2 but without tRNA accounted for almost half of all ribosomes in the dataset (Fig. 2, C and D, and Fig. 3A). This state showed density for SERPINE1 mRNA-binding protein 1 (SERBP1) and thus accounts for hibernating ribosomes (15, 22) (fig. S9G), in which the HHT density was visible (fig. S7C). The ‘P’ state increased in abundance compared to the untreated cells, whereby the ‘A, P’ states showed a similar abundance (Fig. 2C). The HHT molecule was also resolved at the ‘A, P’ state but not in other less populated classes due to the lower resolution (Fig. 3B). We therefore combined the particles from the four remaining less abundant classes. The resulting average also displayed density for HHT (fig. S7C). The positions of A-tRNA and P-tRNA at ‘A, P’ states were similar in untreated and HHT-treated cells (Fig. 3B) (9). One of our classes appeared similar to a potential ‘compact eEF2, A, P’ state (fig. S9D) (23), although the respective local resolution prevents a definitive assignment due to a lack of secondary structure. Finally, two states with Z-tRNA were considerably more abundant compared to untreated cells (Fig. 2D and fig. S9, C and E), showing the typical features of tRNA shape at the Z site (fig. S9, E and H, and movie S3). We conclude that HHT treatment alters the translation landscape in the cellular context in an unanticipated way. It results in the accumulation of ribosome hibernation instead of the ‘A, P’ state, which may be representative of the mechanism of the drug action in cancer therapy.

**Fig. 3.**
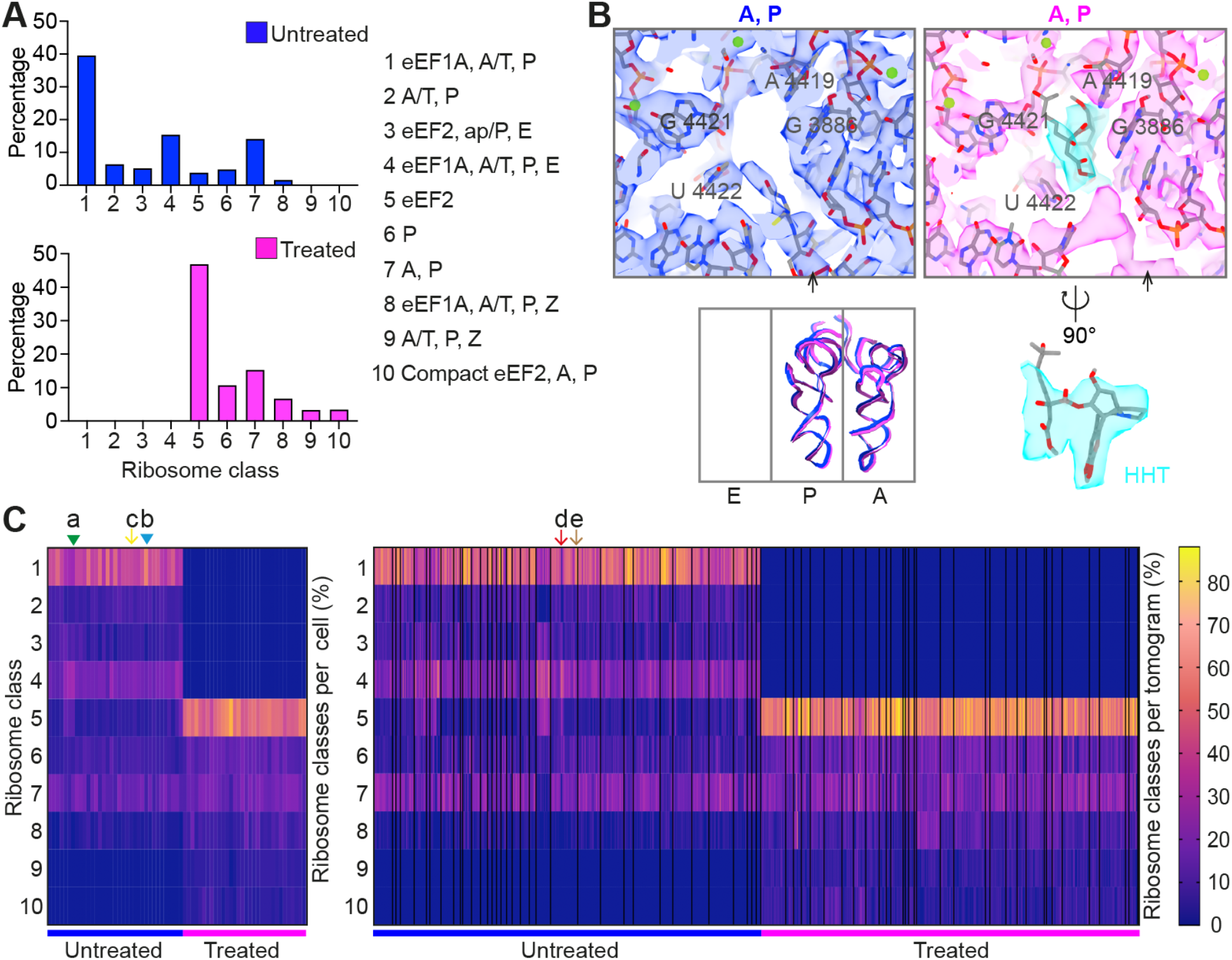
Distribution of ribosome states in cells. (**A**) Percentage of ribosome states from untreated and HHT-treated cells. (**B**) The PTC of ribosomes at the ‘A, P’ state from the two datasets (Materials and methods). Structural overlay of tRNAs at ‘A, P’ states with models determined from previous studies (Materials and methods). Black arrow, P-tRNA. HHT is colored in cyan. (**C**) Heat map of individual ribosome states from the untreated dataset (35 cells, 358 tomograms) and treated dataset (32 cells, 352 tomograms). Each column represents one cell (left) or one tomogram (right). Black lines divide the tomograms from different cells (right heatmap). The class number is the same as in (A). Percentage of ‘eEF1A, A/T, P’ state: 25% in cell a (green triangle, the lowest in the dataset), 56% in cell b (blue triangle, the highest in the dataset), 37% in cell c (yellow arrow), 5% in tomogram d (red arrow), 50% in tomogram e (brown arrow). Tomograms d and e are from the same cell c.

To investigate cell-to-cell variability, we analyzed the distribution of the ribosome states across individual cells that are captured by multiple tomograms. We identified some degree of cell-to-cell variability (fig. S11A), for example, the ‘eEF1A, A/T, P’ state varied from 25% to 56% in untreated cells (Fig. 3C). Overall, the signal observed in multiple tomograms of the same cell was similar, with some notable exceptions that may imply local variability (Fig. 3C). Interestingly, the abundance of the ‘eEF1A, A/T, P’ (class 1) and ‘eEF1A, A/T, P, E’ (class 4) was largely anticorrelated in untreated cells (fig. S11, B and C), while the sum of eEF1A-bound states was relatively consistent, suggesting that its total concentration may be limited (fig. S11D). In contrast, the abundance of ribosomes containing eEF2 was much more diverse between different cells (fig. S11D).

### Polysomes are impaired upon the HHT treatment

We analyzed the spatial organization of different ribosome states within polysomes in treated and untreated cells. The densities accounting for neighboring ribosomes that are commonly observed in the subtomogram averages of untreated cells were reduced in HHT-treated cells (fig. S12A). Furthermore, the ribosome distribution was more dispersed subsequent to treatment (Fig. 4, A and B), both implying that polysomes might be largely abolished. To test this hypothesis, we assigned ribosomes to polysomes based on a distance threshold of 9 nm from the mRNA exit to entry sites of neighboring particles (fig. S12B and table S4, Materials and methods). This threshold grouped 30.2% of all ribosomes into arbitrary polysomes in untreated cells (fig. S12, C to D), which was considerably higher compared to HHT-treated cells (fig. S13, A to E). Most translation states are evenly distributed in monosomes and polysomes, though the ‘eEF2, ap/P, E’ is relatively frequent in polysomes, contrasting the ‘eEF1A, A/T, P, E’ and ‘P’ states (Fig. 4D). The ‘eEF2’ state was prevalent in monosomes (fig. S14A) and has much less neighboring density (fig. S14, C and D), underlining the notion that these ribosomes are hibernating. In polysomes, the ‘eEF2, ap/P, E’ state (class 3, see Fig 3A) was less frequent in the first leading ribosome as compared to the trailing ribosomes (fig. S13F). Classification of all untreated ribosomes with a dedicated mask resolved the low-resolution structure of di-ribosome, in which the mRNA density was visible (fig. S15A) (24, 25). This di-ribosome resembled the ‘top-to-top’ configuration (t-t, the central protuberance of both ribosomes facing a similar direction) (26, 27) (fig. S15B). Two alternative other arrangements of pairs were apparent: one with the central protuberance of the ‘i+1’ ribosome towards down (t-d), the other towards up (t-u) (fig. S15, C and D). Although the abundance of these configurations declined in the treated cells (fig. S15, D and E), the center-to-center and exit-to-entry distance of these pairs were indistinguishable from untreated cells (fig. S15F).

**Fig. 4.**
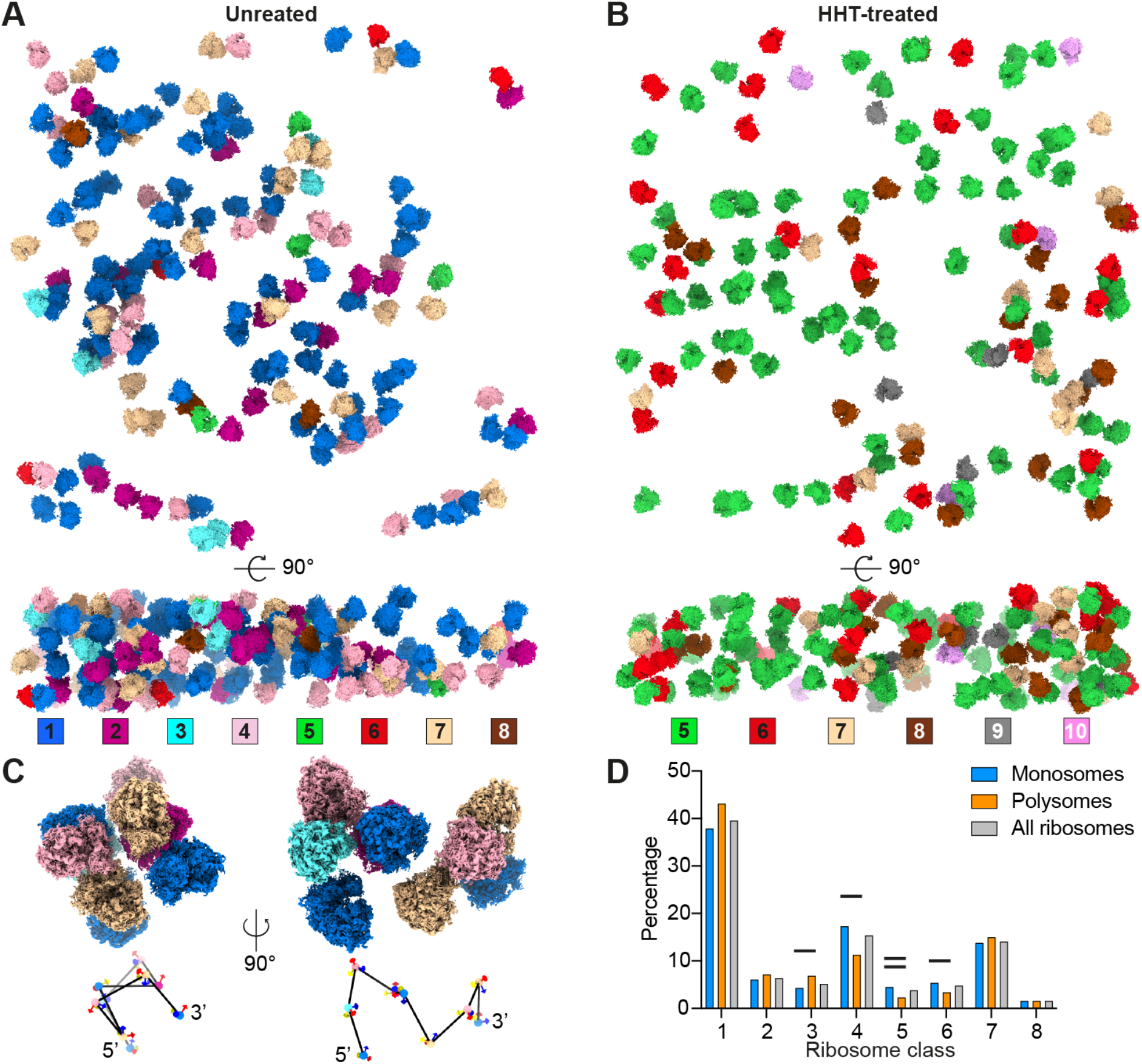
Spatial and functional analysis of Polysomes. (**A and B**) Distribution of ribosome states in a representative tomogram from an untreated (A) and treated cell (B). (**C**) A polysome and the putative mRNA path. The center-to-center distance of the neighboring ribosomes: 27.3 ± 1.8 nm (mean ± SD, n=8). (**D**) Distribution of ribosome states in monosomes, polysomes and all ribosomes from 358 tomograms of untreated cells. The black line above the column illustrates that the abundance of ribosome states in polysomes differs by more than 50% compared to the monosomes. One black line, 50% -95%; two black lines, >95%. Class numbers are the same as in Fig. 3A.

As expected, membrane-associated ribosomes showed a different orientation with the exit tunnel towards the membrane (fig. S16, A to C). The translation state distributions of soluble and membrane-bound ribosomes were similar (fig. S16D). Surprisingly, seven conformations of the expansion segment ES27L were revealed by the classification of membrane-associated ribosomes (fig. S17 and tables S2 and S3). One of which contained a long stretch of ES27L that associates with the ErbB3 receptor-binding protein (Ebp1) on the surface of the 60S and resembles a previously published in vitro structure (28, 29) (fig. S17, A and B). The percentage of Ebp1-associated ribosomes was below 20% in most untreated cells, while the abundance was over 25% in all HHT-treated cells (fig. S18A). Whether Ebp1 is related to specific translation states remained unclear (28, 29). Our analysis revealed that a subfraction of Ebp1-bound ribosomes is observed in all apparent intermediates of the elongation cycle (fig. S18B).

### HHT binds the free 60S associated with eIF6

Finally, we investigated the free 60S and 40S in the cytoplasm of human cells. The free 60S showed Ebp1 and ES27L density in both datasets (Fig. 5, A and B, fig. S19, A and B), which was similar to the Ebp1-associated 80S ribosome. However, the 60S was much more abundant in treated cells than in untreated cells (Fig. 5C). Eukaryotic initiation factor 6 (eIF6) binds the 60S to prevent premature 60S binding with 40S (30, 31). Our data show that the 60S did not contain eIF6 in the cytosol of native untreated cells, contrasting HHT-treated cells (Fig. 5D). These results support the notion that eIF6 prevents the association between large and small ribosomal subunits in cells (*32*–*34*) (Fig. 5, C and D, and fig. S19, C to E). Strikingly, the HHT density was resolved in the 60S from treated cells (Fig. 5E). Thus, we conclude that HHT can bind not only the PTC of different states of 80S ribosome (Fig. 3B and fig. S7C) but also the free 60S decorated with eIF6, which may block the assembly of 40S and 60S in the cytosol of the cell.

**Fig. 5.**
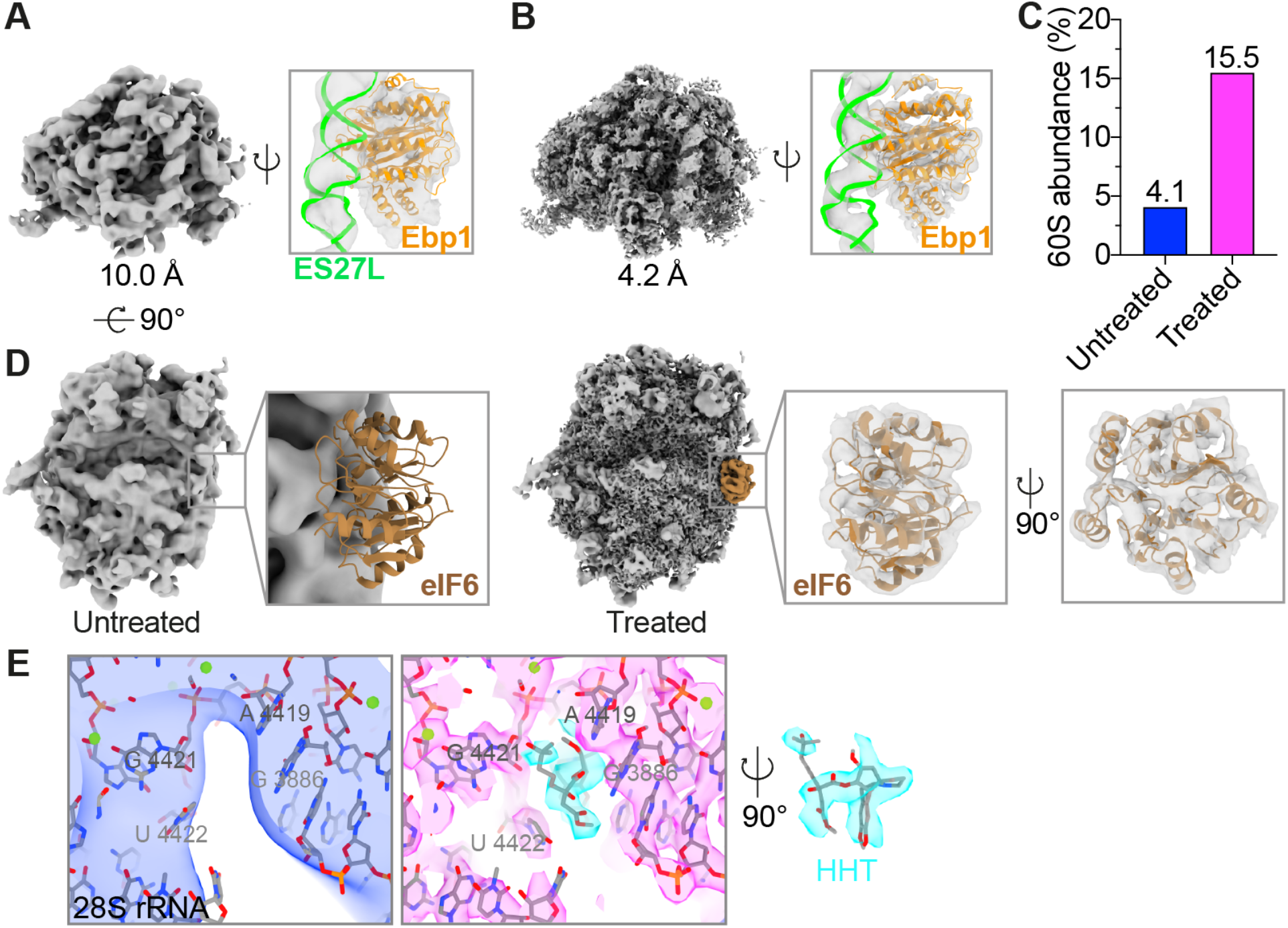
Structure of 60S in human cells. (**A** and **B**) Structures of free 60S in the cytoplasm of untreated cells (A) and HHT-treated cells (B). ES27L and Ebp1 (PDB: 6SXO) are fitted into the 60S maps. Although Ebp1 was not confidently assigned to the 60S from untreated cells due to the low resolution, Ebp1 was fitted better than other factors (amino-terminal acetyltransferases and nascent polypeptide-associated complex) binding the tunnel exit. (**C**) Percentage of the 60S in untreated and HHT-treated cells normalized to the number of 80S ribosomes in the respective dataset. In untreated cells, abundance = 60S/(60S+80S) = 1,693/(1,693+39,402). In treated cells, 60S/(60S+80S) = 7,176/(7,176+39,070). (**D**) The structure of 60S in untreated cells with eIF6 (PDB: 6LSR) fitted into the corresponding position indicates that eIF6 is missing (left). eIF6 fitted into the corresponding density of the HHT-treated 60S (Right). Large subunit GTPase 1 (LSG1), NMD3 and ZNF622 (PDB: 6LSR) are not present in the HHT-treated 60S structure. (**E**) The PTC of free 60S from the two datasets (Materials and methods). HHT is colored in cyan.

## Discussion

Our extensive cryo-ET analysis stresses the feasibility of obtaining structures at near-atomic resolution and visualizes an anti-cancer drug inside human cells. Interestingly, the local resolution in the ribosome core in our study is limited by the pixel size. This illustrates that technical obstacles that hinder high resolution in cellular tomography have been overcome. Improvements in energy filtering (35), image processing (21), and tight control over specimen thickness (36, 37) enable close-to-atomic resolution, as long as the number of particles that can be obtained is sufficient. Further improvements in the automatization of specimen preparation techniques may thus be instrumental in pushing the resolution for other macromolecular assemblies analyzed inside cells (37–42).

Our study provides novel insights into the translation landscape, the coordination of ribosome activity within polysomes and the diverse arrangements of ES27L inside human cells. One may use the same technology in the future to investigate ribosome, translation and mRNA quality control pathways in the context of human cells. Intermediates that have not yet been observed inside cells may be enriched by introducing kinetical bottlenecks, through genetic or pharmaceutical perturbation (5, 43, 44). Our study may set the stage for the analysis of the drug susceptibility of human individuals.

## Acknowledgments

We thank Patrick C. Hoffmann, Maarten Tuijtel and Tomas Majtner from the Department for Molecular Sociology at the Max Planck Institute of Biophysics for advice on membrane segmentation, data collection and data processing. We thank all the members from the Central Electron Microscopy facility and the Department for Molecular Sociology at the Max Planck Institute of Biophysics, Andre Schwarz, Erin Schuman and the Department of Synaptic Plasticity at the Max Planck Institute of Brain Research for support and advice. We acknowledge the support from the Max Planck Computing and Data Facility. We thank Marina Rodnina, Niels Fischer and Stefanie Böehm for critical reading of the manuscript. We acknowledge the support from the Max Planck Computing and Data Facility.

## Author contributions

Conceptualization: H.X., M.B.; Methodology: H.X., B.T., I.K., J.P.K.; Investigation: H.X., R.T., I.K., J.P.K., S.W., B.T.; Visualization: H.X., B.T., M.B.; Funding acquisition: M.B.; Project administration: H.X., M.B.; Supervision: H.X., M.B.; Writing – original draft: H.X., B.T., M.B.; Writing – review & editing: H.X., R.T., I.K., J.P.K., S.W., B.T., M.B.

## Data and materials availability

EM maps have been deposited in the Electron Microscopy Data Bank (EMDB) under accession numbers: EMD-16721, 16725, 16726, 16727, 16728, 16733, 16734, 16735, 16736, 16737, 16738, 16739, 16740, 16741, 16742, 16743, 16722, 16744, 16748, 16747, 16749, 16750, 16751, 16752, 16754, 16755, 16756, 16757. All these maps will be released upon publication. The code for polysome tracing and analysis will be publicly available on GitHub upon publication and is available upon request during the review process.

## Supplementary Materials

Materials and Methods

Figs. S1 to S19

Tables S1 to S4

References (*45*–*63*)

Movies S1 to S3

