## Supplementary_Materials for "Translation dynamics in human cells visualized at high-resolution reveal cancer drug action"

**This PDF file includes:**

Materials and Methods

Figs. S1 to S19

Tables S1 to S4

References

**Other Supplementary Materials for this manuscript include the following:**

Movies S1 to S3

### Materials and Methods

Cell culture

Flp-In T-Rex 293 cells (Invitrogen) were maintained in DMEM (Sigma) supplemented with 10% fetal bovine serum (Gibco) at 37°C with 5% CO2.

Cryo-ET sample preparation

R2/2 gold grids (200 mesh, Quantifoil) were glow discharged using a Pelco easiGlow device for 90 sec on both sides and placed in 3.5 cm cell culture dishes (MatTek). 2 ml cell suspension (175,000 cells/ml) was seeded in the dish containing grids and placed in the incubator. Cells without HHT treatment were cultured for 5 hours (h) before plunge freezing. For HHT (Santa Cruz Biotechnology) treatment, cells were cultured without HHT for 3 h and then treated with HHT at a final concentration of 100µM for 2 h before plunge freezing. For plunge freezing, grids were blotted from the backside for 6 sec using a Leica EM GP2 plunger at 70% humidity and 37°C, plunged into liquid ethane, and stored in grid boxes in liquid nitrogen. Plunge-frozen grids were FIB-milled under cryo-conditions with an Aquilos FIB-SEM (Thermo Fisher Scientific) as previously described (*45*). In brief, grids were coated with an organometallic platinum layer for 15 sec using a gas injection system. Cells were then stepwise milled using decreasing gallium ion-beam currents of 0.5 nA to 30 pA. In total, 35 cells were milled for the untreated sample and 32 for the HHT-treated sample.

Cell viability assay

Cell viability was measured using CellTiter-Glo 2.0 Cell Viability Assay (Promega) according to the manual. In brief, cells at a concentration of 175,000 cells/ml were cultured in 96-well plates. For treated cells, 100 µM HHT was added 3 hours after seeding (fig. S6B). 100µl of CellTiter-Glo. 2.0 Reagent was added at the corresponding time point (fig. S6B) and mixed for 2 minutes at room temperature. The luminescent signal was measured using a microplate spectrofluorometric reader (TECAN SPARK). Representative data from three repeated independent experiments are shown as mean ± standard deviation (SD). The statistical analysis was performed using two-tailed unpaired *t*-tests in GraphPad Prism. Previous work indicates that the impact of HHT on cell viability depends on the cell type, detection method and HHT treatment time: Beas-2b cells, 18 h, cytotoxicity concentration 50% ($CC_{50}$) > 10 µM (*46*); Huh7 cells, 48 h, $CC_{50}$ = 0.0218 µM (*47*); Vero E6 cells, 48 h, $CC_{50}$ = 59.75 µM (*48*); U937 cells, 48 h, $CC_{50}$ > 0.1 µM (*49*). This makes CC50 less reliable than naturally expected (*50*).

<u>BCA assay</u>

The protein concentration was assessed with a bicinchoninic acid (BCA) assay kit (Thermo Fisher Scientific), following the standard protocol. For the untreated sample, cells were cultured in the 24-well plates for 5 h before measuring the protein concentration. For treated cells, HHT was added 3 h after seeding at a final concentration of 100µM. 2 h later, cells were lysed and the protein concentration was measured. Representative data from three repeated independent experiments are shown as mean ± standard deviation (SD). The statistical analysis was performed using two-tailed unpaired *t*-tests in GraphPad Prism.

<u>Data acquisition</u>

Tilt series were acquired on a Titan Krios G4 (Thermo Fisher Scientific) operated at 300kV and equipped with Selectris X imaging filter and Falcon 4 direct electron detector, at 4K x 4K pixel dimensions, pixel size of 1.223 Å, tilt range of -60° to 60°, 2° tilt increment, target defocus of -1.5 to -4.5 µm and a total dose of 120 to 150 e/Å$^2$ per tilt series, using SerialEM software (*51*). Target areas were selected in the cytoplasmic region of the cell (fig. S1A).

<u>Tomogram reconstruction and 80S ribosome template matching</u>

Tilt series were aligned with patch-tracking in IMOD and reconstructed as back-projected tomograms with SIRT-like filtering of 10 iterations at bin4 (*52, 53*). The alignment files were used for tomogram reconstruction in Warp 1.0.9 (*21*). In total, 358 tomograms were analyzed from untreated cells and 352 from HHT-treated cells. Template matching was performed with STOPGAP (*54*). For this, 1,080 potential 80S ribosomes from the untreated dataset and 1,031 from the HHT-treated dataset were manually picked and refined in RELION 3.1 as templates. The template matching results were visually checked in napari (*55*) (movie S1). To utilize all potential 80S ribosomes in the datasets, the coordinates with the top 800 constrained-cross-correlation (CCC) values in each tomogram were extracted as sub-tomograms in Warp. In total, 286,400 untreated ribosome sub-tomograms and 281,600 HHT-treated were reconstructed.

Refinement and model building

Sub-tomograms from template matching were classified and refined in RELION 3.1 (*56*). 39,402 particles from the untreated dataset and 39,070 from the treated dataset were assigned to the 80S ribosome class. Three repeats were performed with different initial references for comparison (figs. S3A and S8A). After refinement in RELION, multi-particle refinement of tilt series was performed in M 1.0.9 as previously described (*21*). Refinement of geometric and CTF parameters was performed sequentially. Fourier shell correlation (FSC) calculation and local resolution estimation were conducted in RELION and M. The untreated and treated 80S ribosomes structures reached ~3.2 Å resolution after the last refinement iteration in M. The atomic models of the human ribosome (PDB: 6XA1 and 6QZP) (*12*, *57*) were used as initial models and fitted into the 3.2-Å untreated and HHT-treated ribosome maps, followed by PHENIX real-space refinement (*58*) and adjustment in Coot (*59*). For visualization, the two above refined models were rigid-body-fitted into the untreated and HHT-treated maps in Figs. 3B and 5E, and fig. S7C. Density maps and molecular models were visualized using ChimeraX (*60*).

Subtomogram classification

To classify translation states, a tRNA mask and factor mask (fig. S2 and S7) with soft edge covering the tRNA path (A, P and E sites) and elongation factor binding sites were used sequentially in RELION 3.1 (*5*). First, 39,402 untreated and 39,070 HHT-treated ribosomes identified from the two datasets were classified with a tRNA mask (10 classes, T = 4, 35 iterations). Second, classification with a factor mask was performed using the following parameters: 3 or 5 classes, T = 4 or 5, 35 iterations. The ribosome classes with fewer than 500 particles were discarded after each round of classification because they could not be unambiguously assigned to specific states due to the low resolution. Five repeats were carried out to validate the classification with the same parameters mentioned above (figs. S3B and S8B). Finally, eight classes were identified in untreated cells and six classes were classified in treated cells. tRNAs (A, P, E and Z), eEF1A and eEF2 atomic models were obtained from PDB (6Z6M, 5LZS, 6MTB, 6TNU) and fitted into our density map of each state (*4*, *13–15*).

An elliptic mask covering the potential ES27L and membrane of the ribosome was used to classify the ribosomes with membrane, ES27L and Ebp1 (T = 4, 40 iterations). PDB 6SXO was rigid-body-fitted into our ribosome structures with Ebp1 (*28*). 654 membrane-bound ribosomes were identified. For validation, membrane-associated ribosomes were counted manually in tomograms containing membranes, showing a similar number as the classification. The results were also checked by mapping the identified membrane-bound ribosomes back into tomograms with ArtiaX in ChimeraX (*60*, *61*). A representative tomogram with membranes was segmented using Amira-Avizo 2021.1 (Thermo Fisher Scientific). The number of membrane-bound translation states is calculated by analyzing the shared particle coordinates between the membrane-associated ribosomes and individual translation states. First, the membrane-bound ribosome classification was carried out independently of the classification of the translation states. Then, each translation state was assigned to the membrane-associated ribosomes.

To classify the di-ribosome from the 39,402 untreated ribosomes, sub-tomograms were extracted with a bigger box size that could accommodate three ribosomes. After refinement in RELION 3.1, the classification was performed with a sphere mask focusing on the trailing ribosome (i+1). PDB 6I7O and 7QVP were fitted into our map (*24*, *25*) (fig. S15A).

Ribosome states analysis in individual tomograms and cells

After classification and refinement, 80S ribosome states were mapped back into their original tomograms for calculating their abundance. The tomograms were mapped back to the cells where they come from to represent the abundance of each cell's abundance of ribosome states. We analyzed 358 tomograms from 35 untreated cells and 352 tomograms from 32 treated cells. The heatmaps were prepared in GraphPad Prism. The single-cell clustering analysis was done in Matlab2019b with the clustergram function.

Polysome analysis

The refined positions obtained during the subtomogram averaging were used to trace the polysomes within tomograms. Since the coordinates correspond to the center of ribosomes, they were shifted to entry and exit points (see fig. S12B), resulting in two different sets of coordinates for each ribosome. For each tomogram, the polysome chains were traced using the routine described in table S4. The decision to append a ribosome to a chain was based solely on the distance between the exit point of the leading ribosome (denoted i) and the entry point of the trailing one (denoted i+1), i.e., the rotation of the trailing ribosome with respect to the leading one was not considered by the script. The ribosome with the shortest Euclidean distance (the nearest neighbor) to the leading ribosome was chosen to be the trailing one if it was within the allowed distance threshold. The distances from 2 to 25 nm were tested to determine the optimal threshold (figs. S12C and S13A). For each distance, the threshold was within the ±0.5 nm range (i.e., for 2 nm, the allowed distance was from 1.5 nm to 2.5 nm). The visual inspection of polysomes created with distances between 7 and 12 nm resulted in the threshold of 9 nm. The analysis of polysomes presented in this work corresponds to the distance threshold range from 0 to 9 nm. The polysome chain tracing was implemented in Python and will be publicly available on GitHub, together with Jupyter notebooks that were used to produce presented results.

Angle analysis of ribosome pairs

The distances and orientations between the neighboring ribosomes within polysomes were analyzed similarly to the previous study (*62*). Let $(\varphi, \theta, \psi)_i$ denote Euler angles describing the rotation of the reference to the leading ribosome *i* and let $(x, y, z)_i$ denote its coordinates. To analyze the angular relationships within the neighboring ribosomes, the leading ribosome *i* was rotated to the reference position (called zero rotation) by applying inverse rotation, i.e. it was rotated by $(-\psi, -\theta, -\varphi)_i$. The trailing ribosome i+1 was rotated by its rotation $(\varphi, \theta, \psi)_{i+1}$, followed by a rotation of $(-\psi, -\theta, -\varphi)_i$. This brings the ribosome pair into a common rotation frame (zero rotation of the leading ribosome) while keeping their original angular relationship. The relative orientations of the ribosomal pairs within polysomes are shown in figs. S13D and S15D. For the distance analysis, the exit site coordinates of a leading ribosome were subtracted from both the leading ribosome's exit site coordinates (setting it to zero) and its trailing ribosome's entry site coordinates. The new entry site coordinates of the trailing ribosome were rotated by $(-\psi, -\theta, -\varphi)_i$ to show their position with respect to the zero rotation of the leading ribosome (fig. S13C).

Visual analysis of the neighboring pairs revealed three abundant orientations (fig. S15, B and C). Five representative pairs from each observed group were analyzed to determine their angular relationship. The top-top (t-t) orientation corresponds to the angular difference (*63*) of roughly 90 degrees. The top-down (t-d) and top-up (t-u) have both angular differences of around 115 degrees. The t-d and t-u pairs can be further distinguished by the difference in their φ angles – for t-d the difference is above 100 degrees and for t-u it is below 0 degrees (the Euler angles are always expressed in their canonical form, which yields range ±180 degrees for φ and ψ, and 0 – 180 degrees for θ). Applying these restrictions (with the tolerance of ±10 degrees) onto neighboring pairs within polysomes yielded the following distribution (fig. S15, D and E): 40.9% (t-t), 16.2% (t-d) and 4.5% (t-u) in the untreated dataset and 34.8% (t-t), 0.5% (t-d) and 9.5% (t-u) for the treated dataset.

Free 60S and 40S template matching, classification and refinement

We processed the 60S and 40S similarly to the 80S ribosome (see above). After template matching of free 60S and 40S, several rounds of classification (3 or 4 classes, T = 1, 30 iterations) were performed in RELION 3.1 (*5*). 1,693 60S and 1,682 40S were classified in untreated cells. 7,176 60S and 3,895 40S were classified in HHT-treated cells. The refinement was performed in RELION3.1 and M 1.0.9.

### Supplementary figures

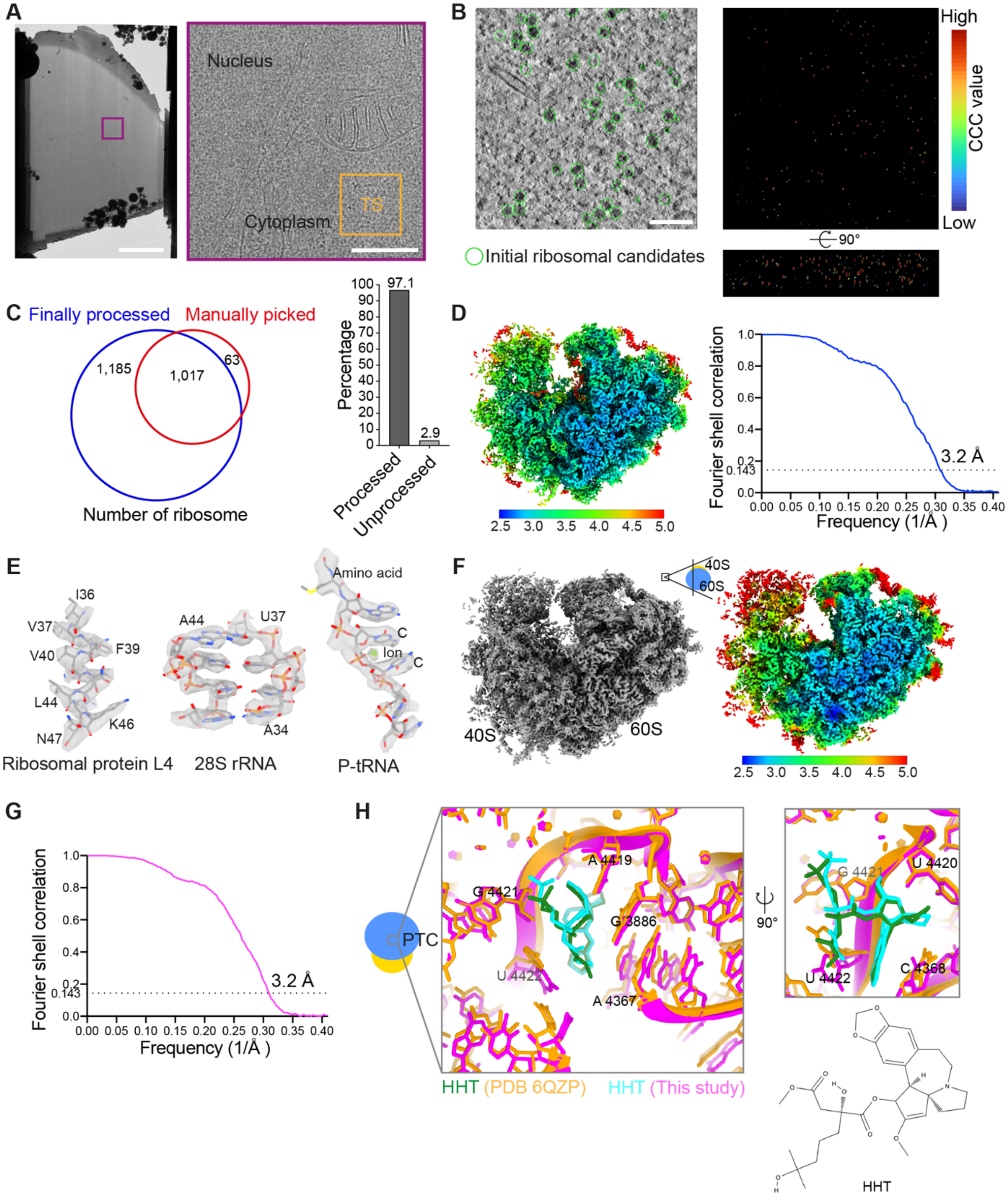


**Fig. S1. Untreated and HHT-treated ribosome maps in human cells.**

(**A**) Transmission electron micrographs of FIB-milled lamellae. Scale bar, 4 μm (left panel); 500 nm (right panel). The orange square indicates a representative region used for data collection. TS denotes tilt series. (**B**) A tomogram slice from an untreated cell. Top 800 potential ribosomes (green circles) per tomogram were selected based on the constrained cross-correlation (CCC) value of template matching for the following classification (left). CCC values from the template matching (right, see also movie S1). (**C**) Evaluation of actually processed and unprocessed ribosomes in this study. 1,080 potential ribosomes (red cycle) are manually picked based on experience from 10 tomograms. In the same 10 tomograms, 2,202 ribosomes (blue cycle) are actually processed using the pipeline (see fig. S2). 63 out of 2,202 (2.9%) potential ribosomes are not processed (right panel). (**D**) 80S ribosome determined from 39,402 particles from untreated cells and the map colored by local resolution. The core regions of the 60S are resolved at 2.45 Å resolution. The Nyquist limit of the dataset is 2.45 Å. The Fourier shell correlation (FSC) curve of the untreated ribosome reveals an overall resolution of ~3.2 Å using the 0.143 criterion. (**E**) Structures of ribosomal protein L4, 28S rRNA and P-tRNA from untreated cells. Green sphere, potential $Mg^{2+}$. (**F**) The structure of the ribosome determined from 39,070 particles under HHT treatment. The map is colored by local resolution. 40S was less resolved possibly owing to the heterogeneity of particles (see Fig. 2, half population of the treated ribosome with unrotated small subunit and half with rotated). (**G**) The FSC curve for the HHT-treated ribosome. (**H**) Structural overlay of HHT-bound ribosome reconstituted in vitro (orange, PDB 6QZP) and the ribosome in this study (Magenta). The chemical structure of HHT (bottom).

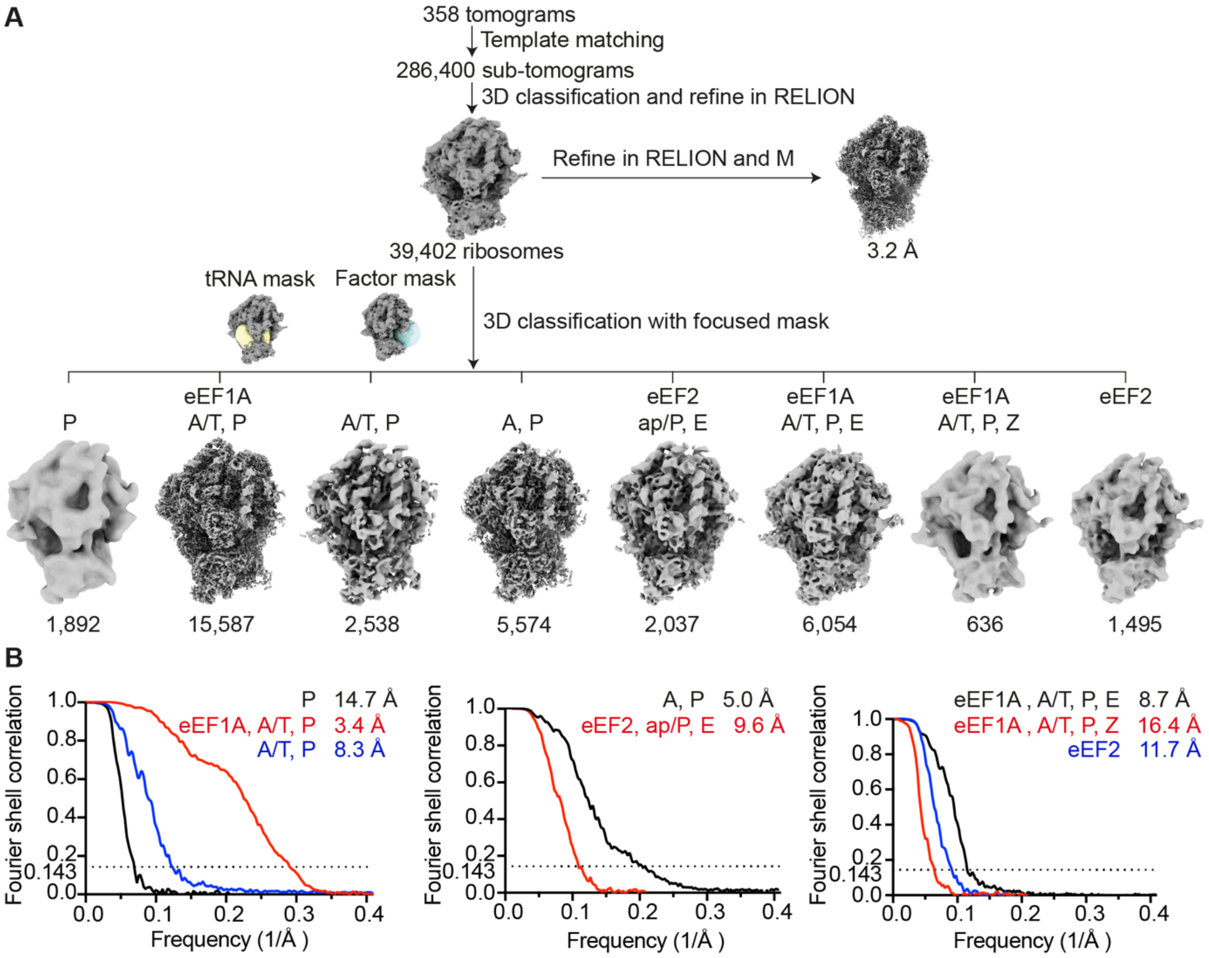


**Fig. S2. Image-processing workflow for the dataset of untreated cells.**

(**A**) Diagram of the cryo-ET data analysis workflow of ribosomes in untreated cells. Template matching in STOPGAP generates 286,400 ribosome candidates. Classification in RELION identified 39,402 ribosomes after removing false positive particles. Classified ribosomes are refined in RELION and M. A focused classification is performed with a tRNA mask covering the tRNA path and a factor mask focusing on the elongation factor binding area. Finally, eight ribosome states are determined. (**B**) FSC curves of the corresponding ribosome structures and the resolution are provided (FSC = 0.143).

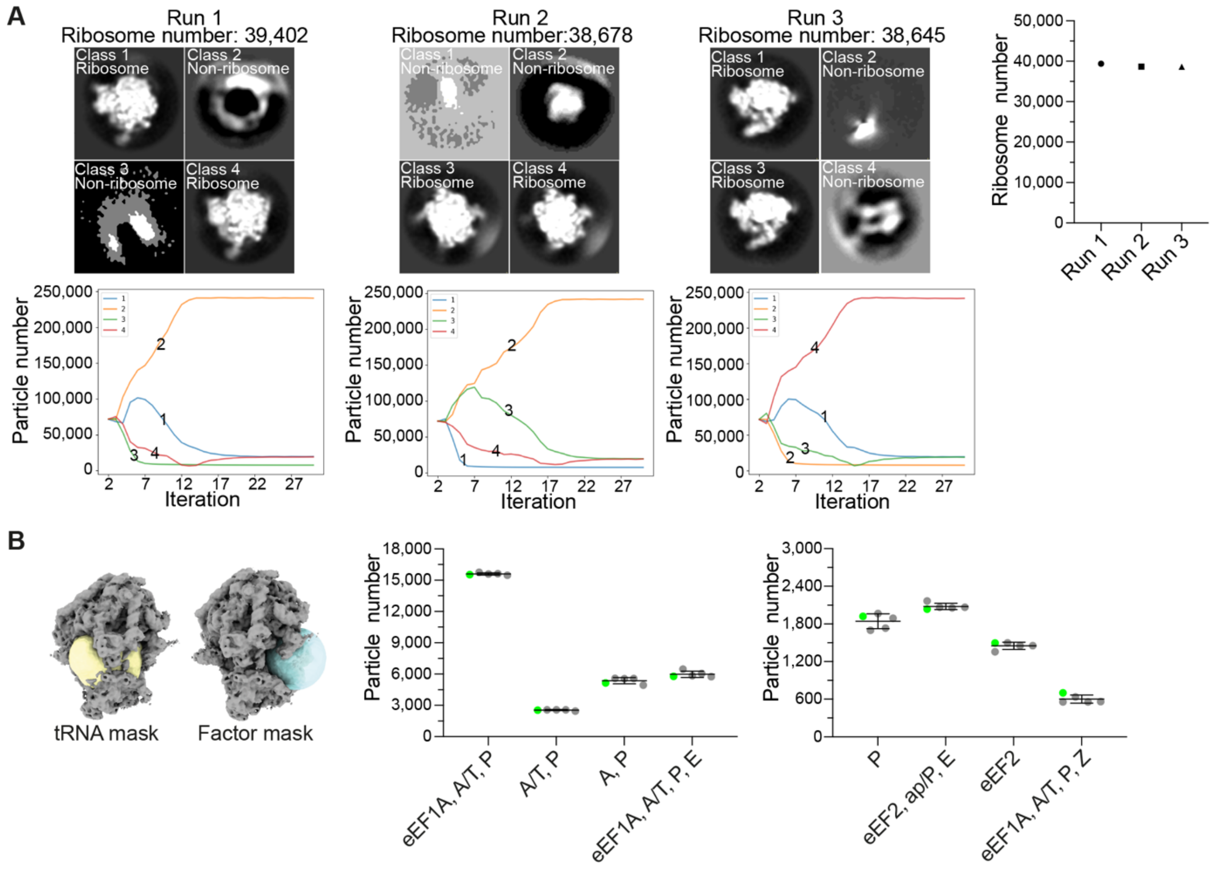


**Fig. S3. Validation of ribosome classification in native untreated cells.**

(**A**) Outputs of three runs to classify ribosomes using different initial references after template matching in untreated cells. In each run, the total number of the classified ribosome is summarized (top right). The curves show the change in the particle number of each class over 30 iterations (bottom panel). The first run is used for the following processing. (**B**) Five repeats of the focused classification with the indicated masks using the same parameters. The repeat marked with green dots is used to calculate the percentage in Fig. 2 and the final refinement in RELION and M.

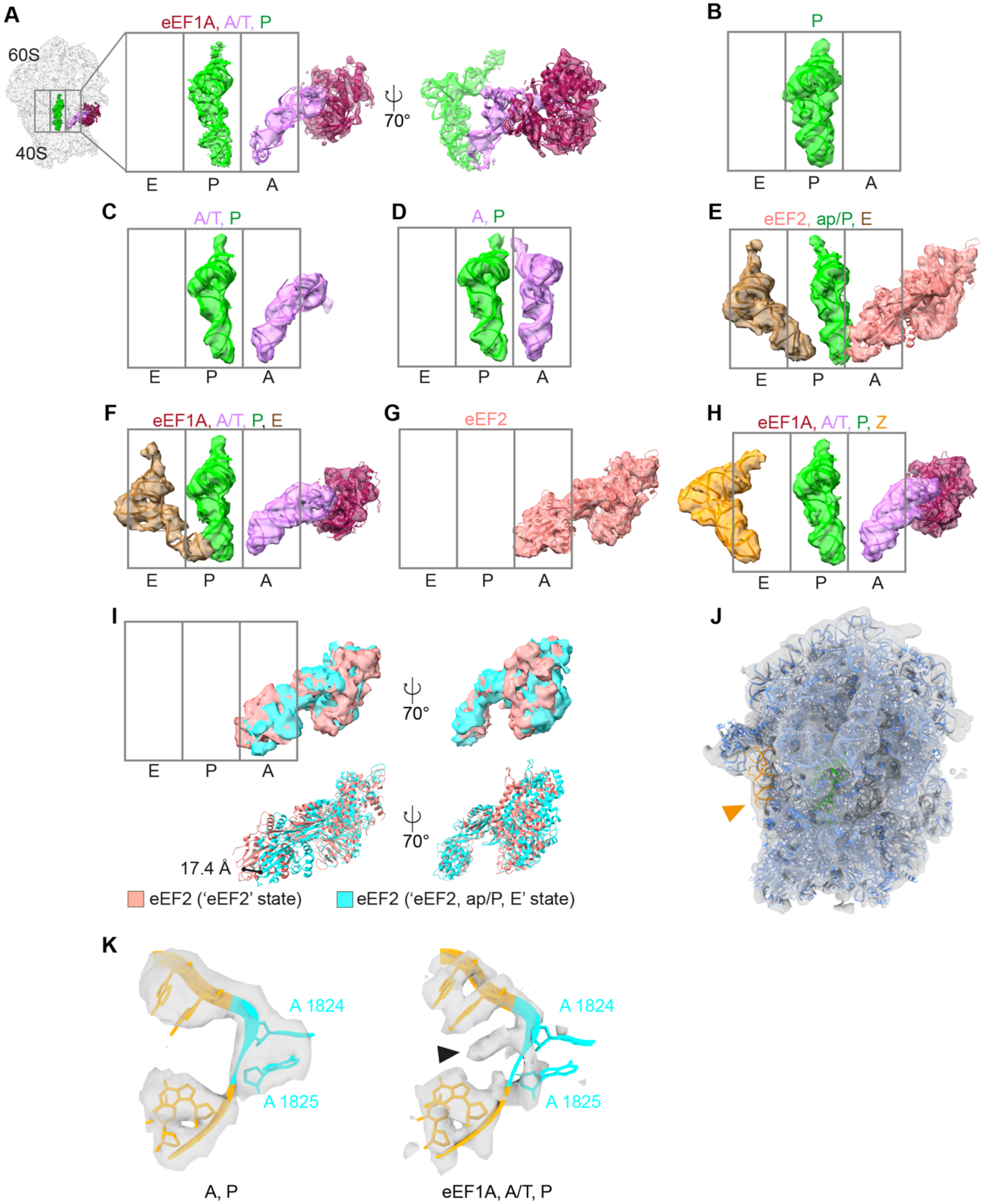


**Fig. S4. Atomic models of tRNAs and elongation factors are fitted into the map of the untreated ribosome.**

(**A** to **H**) The densities of tRNAs, eEF1A and eEF2 from eight ribosome states are rigid-body-fitted with the previously determined atomic model (Materials and methods). The fitted models represent the averaged positions of the nearby tRNA and the elongation factor. Colors: eEF1A (maroon), eEF2 (salmon), A and A/T (lavender), P and ap/P (green), E (brown), Z (tangerine). (**I**) The difference in eEF2 position between the 'eEF2, ap/P, E' state (cyan) and the 'eEF2' state (salmon). The 80S ribosome maps of the two states are fitted first, and then the eEF2 model from PDB 6Z6M is fitted into the corresponding density map. An amino acid in the same region is selected for measuring the distance. (**J**) The atomic model of the 'P, Z' state ribosome (PDB: 6MTB) is fitted into the map of the 'eEF1A, A/T, P, Z' state from untreated cells. Tangerine, Z-tRNA. (**K**) The atomic model (PDB 5LZS) of 18S rRNA (A1822 to A1827) is rigid-body-fitted into the corresponding density of the 80S at 'A, P' and 'eEF1A, A/T, P' states. The decoding nucleotides A1824 and A1825 in the atomic model are in the flipped-out configuration. Triangle points to the potential A1824 density in our EM map.

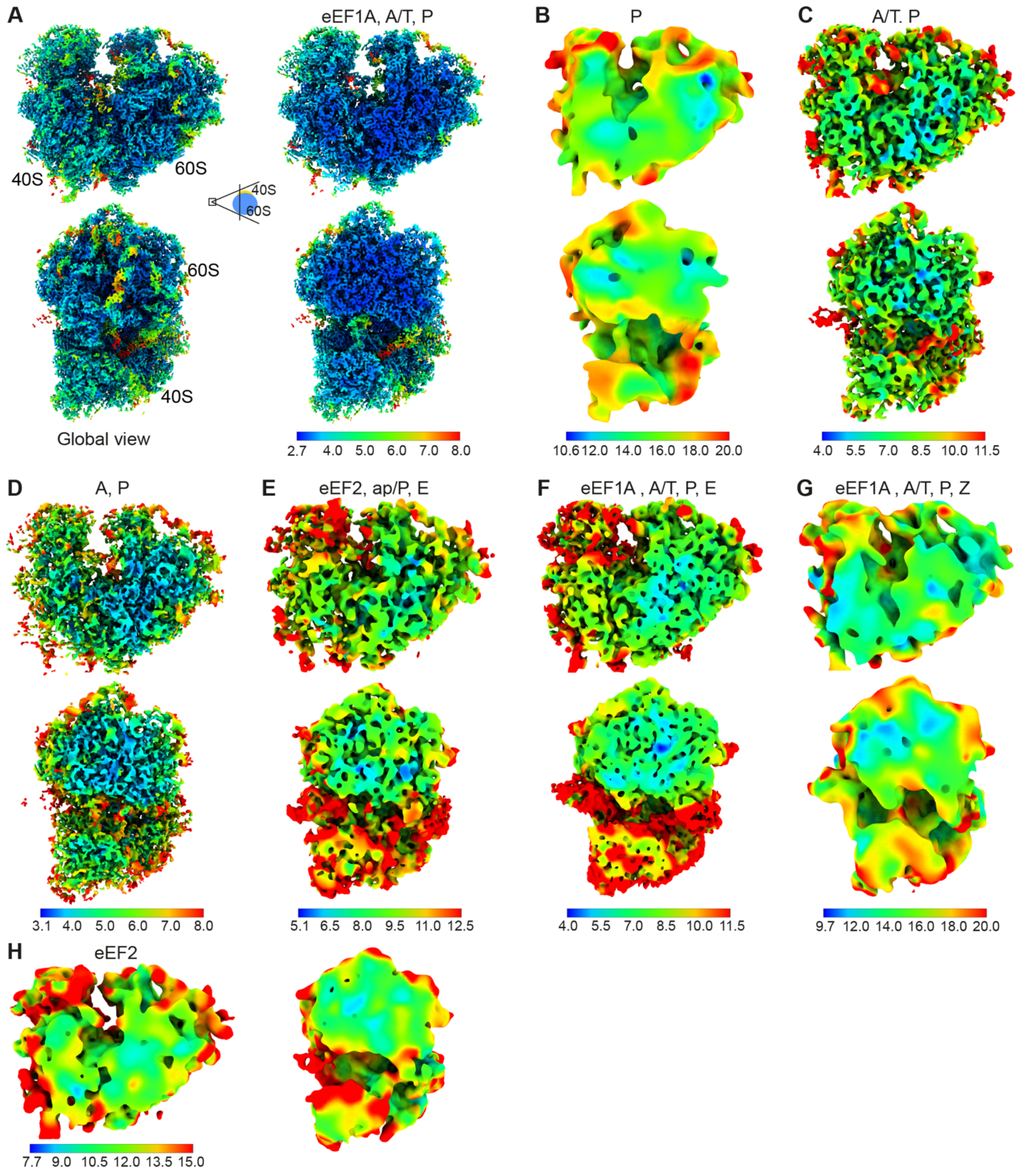


**Fig. S5. Local resolution maps for ribosome classes in the untreated dataset.**

(**A** to **H**) Eight ribosome states identified in the untreated dataset displayed as color-coded local resolution maps calculated in M. Color keys are shown below each panel.

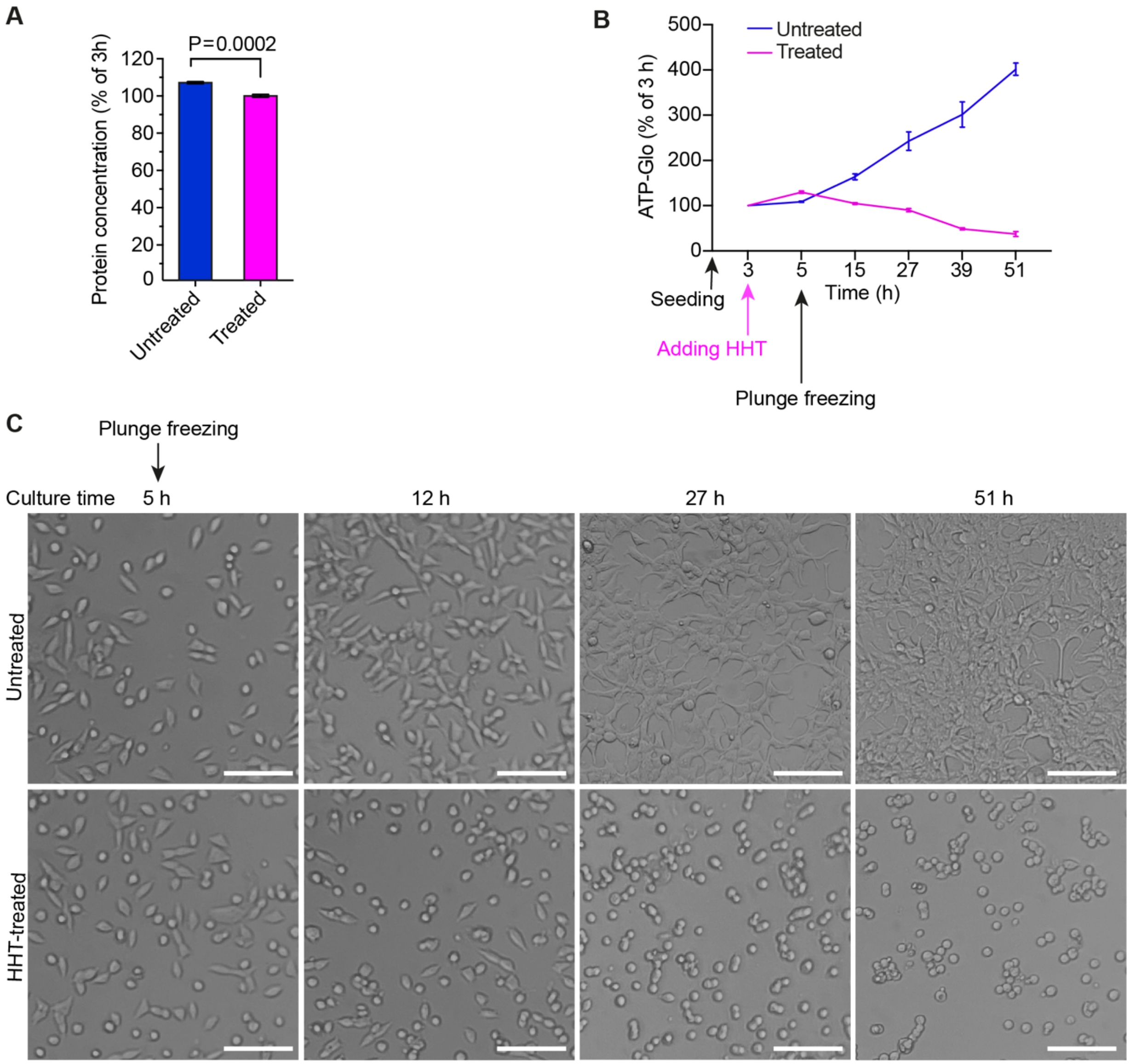


**Fig. S6. The impact of HHT on protein expression and cell viability.**

(**A**) Relative protein concentration 5 h after seeding (the same time point when we did plunge freezing) in untreated and HHT-treated cells. For treated cells, 100 µM HHT was added 3 h after seeding. The protein concentration at 3 h after seeding was set to 100%. The data represent the mean ± SD of three independent experiments. Statistical analysis was performed using two-tailed unpaired *t*-tests in GraphPad Prism. Significantly different ($P < 0.05$). (**B**) Cell viability was assessed by the ATP level in the cells at corresponding time points. For treated cells, 100 µM HHT was added 3 h after seeding. The ATP signal at 3 h after seeding was set to 100%. The data represent the mean ± SD of three independent experiments. (**C**) Cell morphology at different time points with or without HHT treatment. The top panel shows untreated cells 5 h to 51h after seeding as control. The bottom panel shows cells treated with 100 μM HHT. The drug was added 3 h after seeding, meaning the drug treatment time is 2 h, 9 h, 24 h or 48 h from left to right. Scale bar, 100 μm.

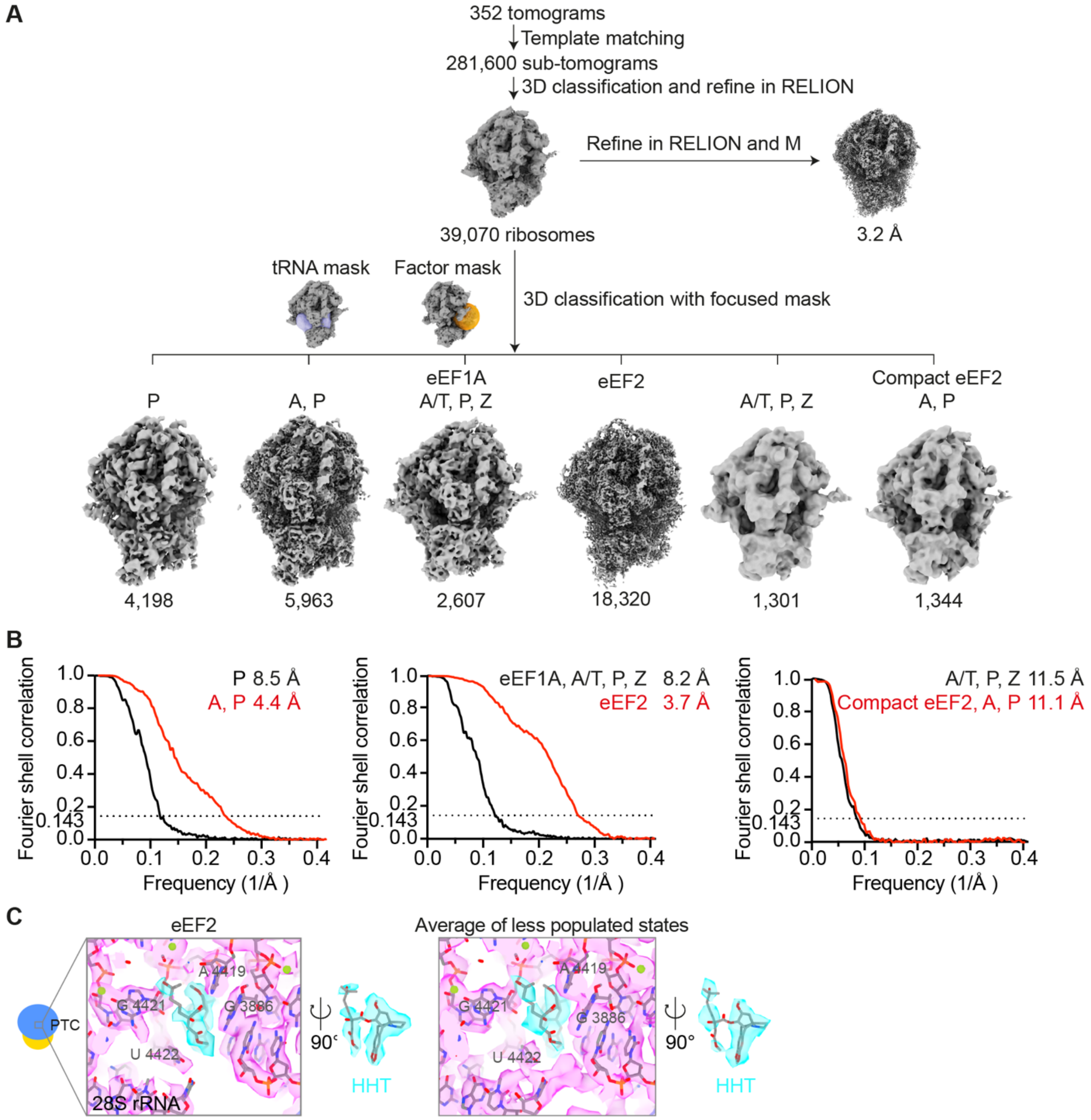


**Fig. S7. Data-processing workflow for the HHT-treated cells.**

(**A**) Diagram of the cryo-ET data analysis workflow of ribosomes in HHT-treated cells. Template matching in STOPGAP generates 281,600 ribosome candidates. Classification in RELION identified 39,070 ribosomes after removing false positive particles. Classified ribosomes are refined in RELION and M. A focused classification is performed with a tRNA mask covering the tRNA path and a factor mask focusing on the elongation factor binding area. Six ribosome states are determined. (**B**) FSC curves of corresponding ribosome states and the resolution are provided (FSC = 0.143). (**C**) The peptidyl transferase center (PTC) of ribosomes from HHT-treated cells. The HHT structure of the 'eEF2' state (left panel). The HHT structure of the 'A, P' state is shown in Fig. 3B. The average map of the four less abundant classes (right panel): 'P', 'eEF1A, A/T, P, Z', 'A/T, P, Z' and 'Compact eEF2, A, P'. HHT is colored in cyan.

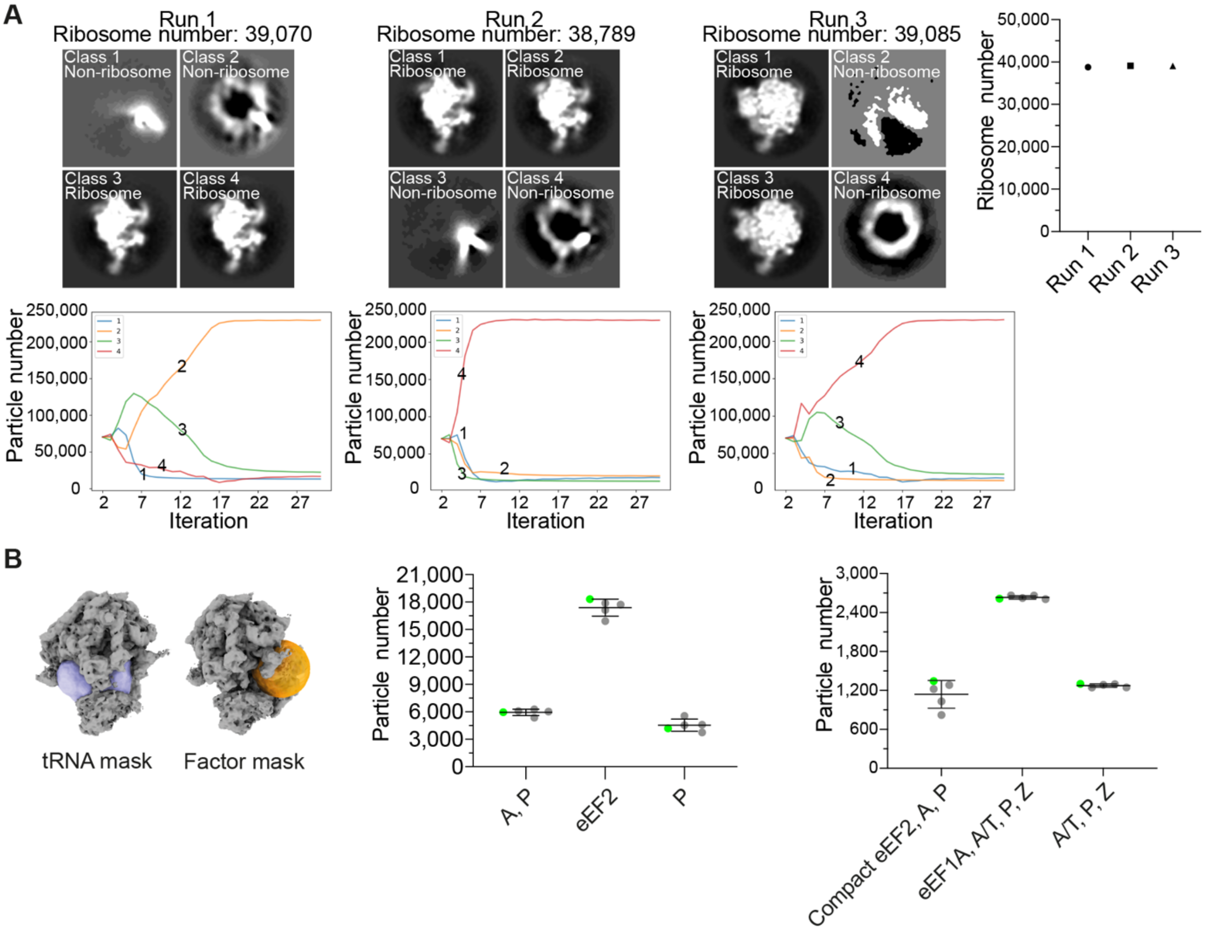


**Fig. S8. Validation of ribosome classification from HHT-treated cells.**

(**A**) Output of three runs of classification to identify ribosomes after template matching in HHT-treated cells. The total number of the classified ribosomes is shown in the top-right panel. The curves show the change in particle number of each class over 30 iterations (bottom panel). The first run is used for the following classifications. (**B**) Five repeats of the classification with masks focusing on the tRNA path and elongation factor binding site. One of the repeats marked with green dots is used for the final refinement in M and for calculating the percentage in Fig. 2.

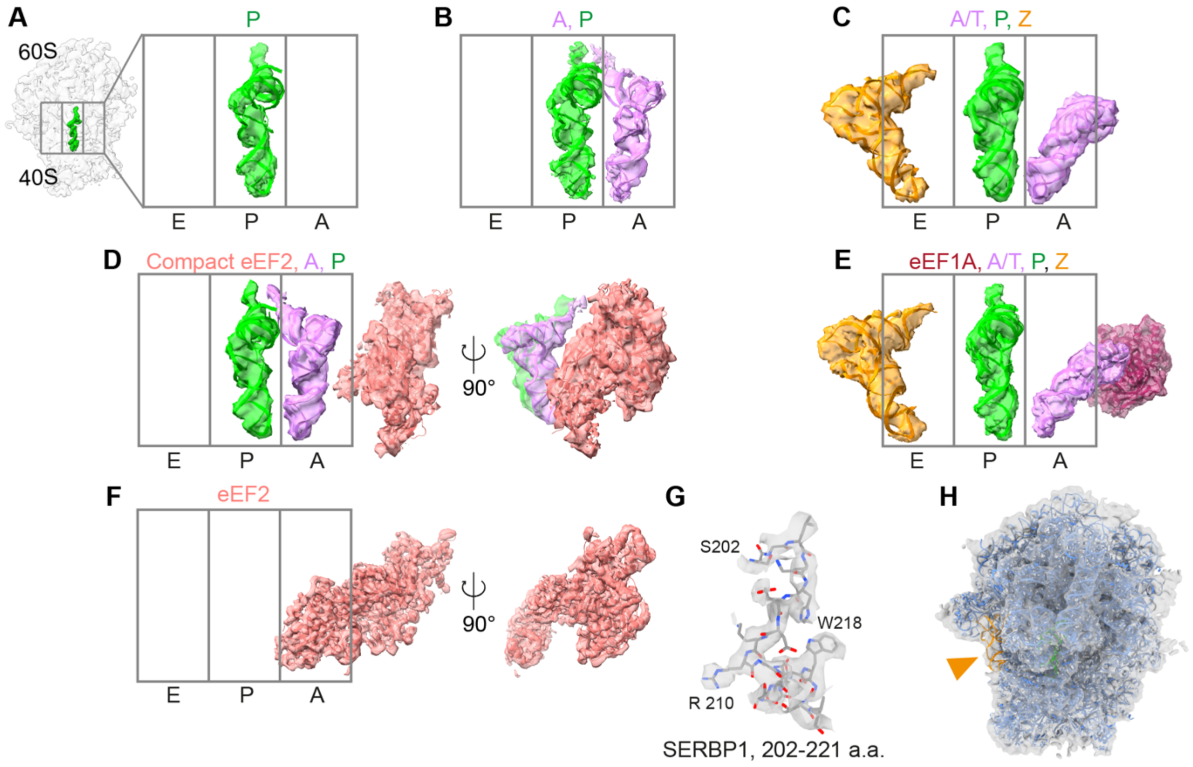


**Fig. S9. Atomic models of tRNAs and elongation factors are fitted into the corresponding densities of the treated ribosomes.**

(**A** to **F**) The densities of tRNAs and elongation factors are fitted with the previously determined atomic model (Materials and methods). Colors: eEF1A (maroon), eEF2 and compact eEF2 (salmon), A and A/T (lavender), P and ap/P (green), E (brown), Z (tangerine). In (D), five individual domains of eEF2 (PDB: 6Z6M) are rigid-body-fitted into the density of compact eEF2. (**G**) The atomic model of SERBP1 (202-221 amino acids) from PDB 6Z6M is fitted in the 'eEF2' map. No densities of eIF5A and coiled-coil domain containing short open reading frame 124 (CCDC124) are found in the 'eEF2' state. (**H**) PDB 6MTB is fitted into our 'eEF1A, A/T, P, Z' state ribosome (see also movie S3). Tangerine, Z-tRNA.

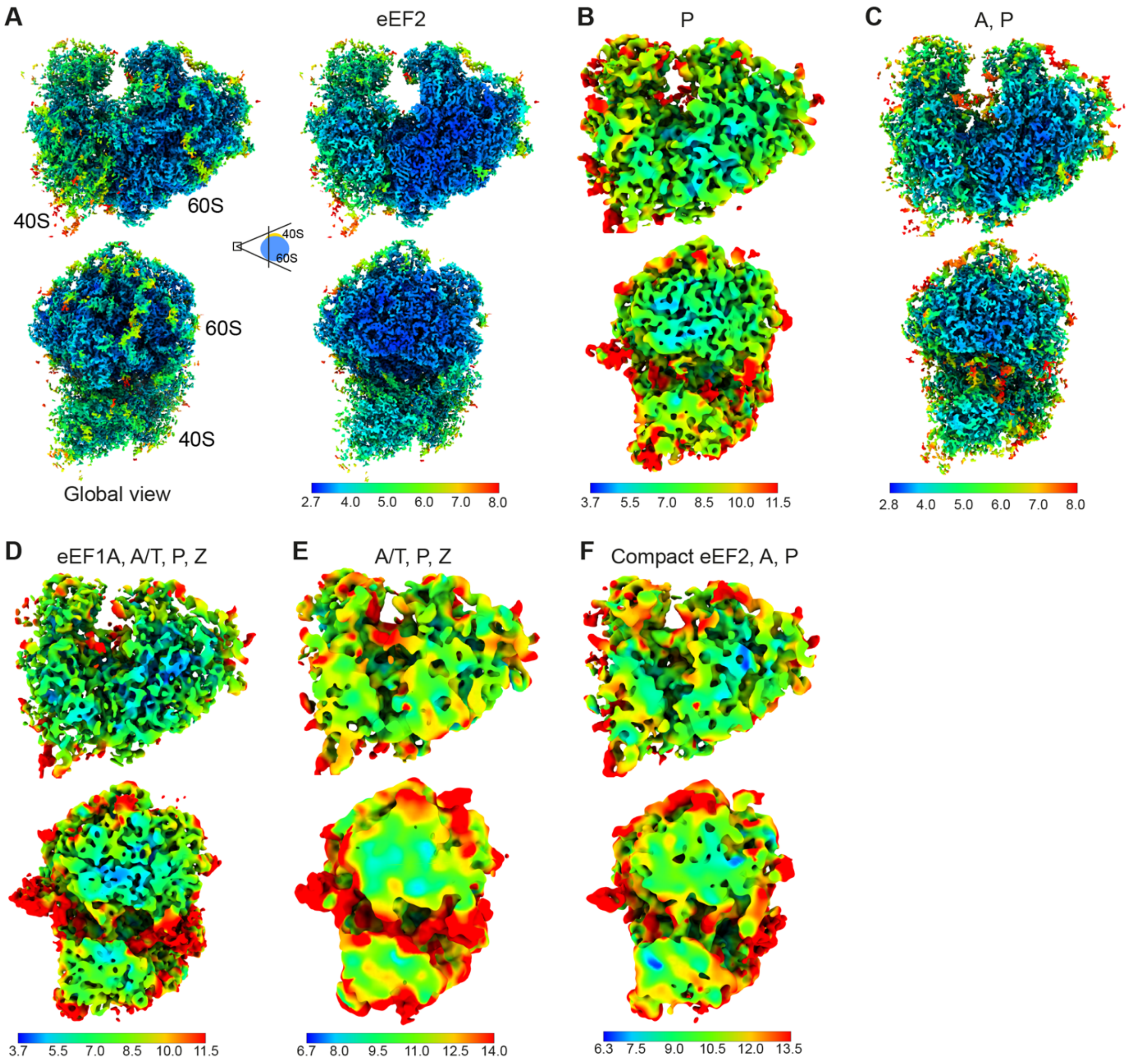


**Fig. S10. Local resolution maps for ribosome classes from the treated cells.**

(**A** to **G**) Six ribosome states are colored by local resolution calculated in M. Color keys are shown in the bottom.

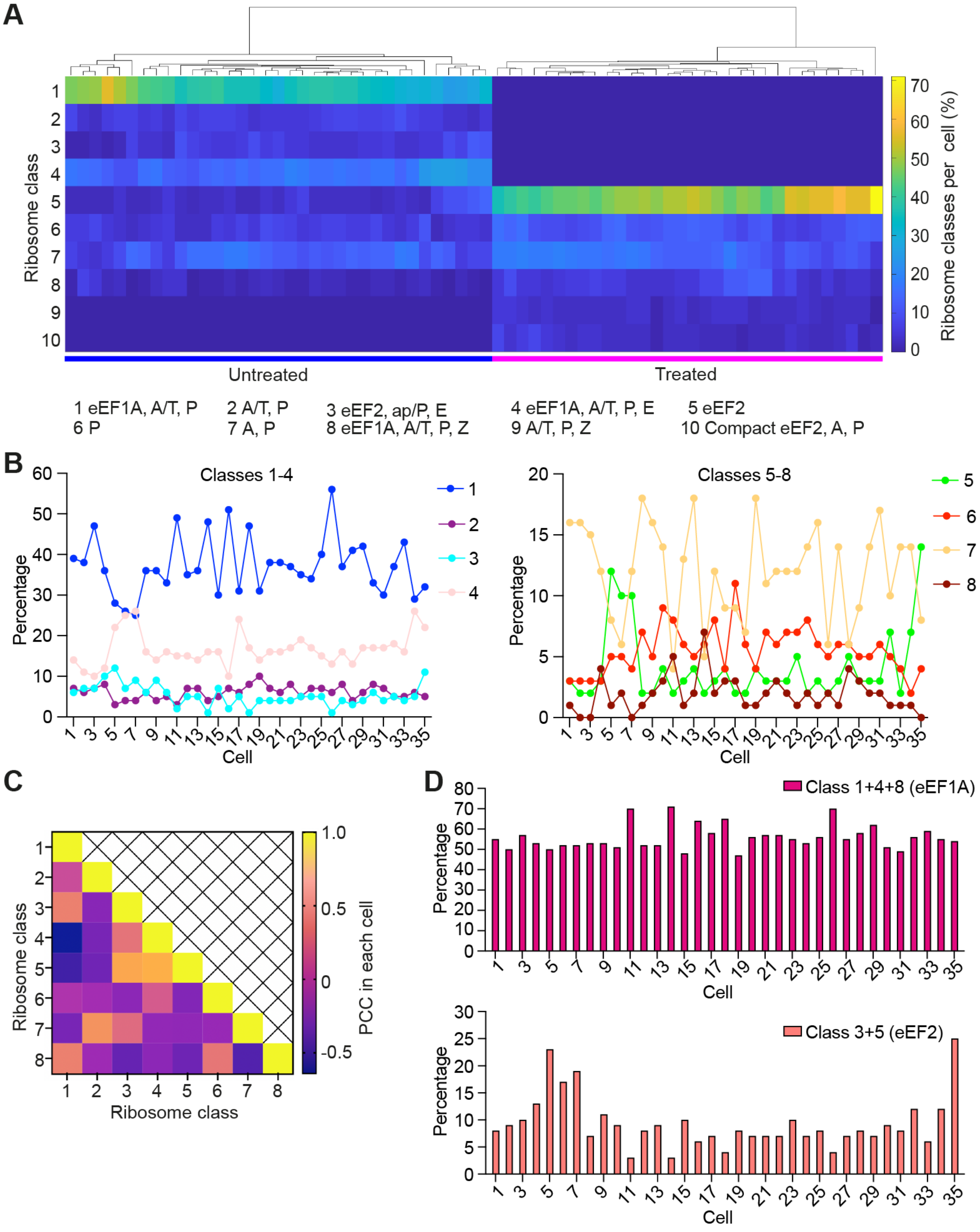


**Fig. S11. Analysis of ribosome states in individual cells.**

(**A**) Clustering analysis of ribosome states in 35 untreated and 32 HHT-treated cells. Each column represents one cell. The analysis was done using clustergram in MATLAB 2019b. (**B**) The abundance of eight ribosome states in individual untreated cells. (**C**) Pearson's correlation coefficient of ribosome states in the same cell (35 untreated cells). The input is the corresponding percentage in (B). The class number is the same as in (A). (**D**) Percentage of ribosome states containing eEF1A or eEF2 in individual cells.

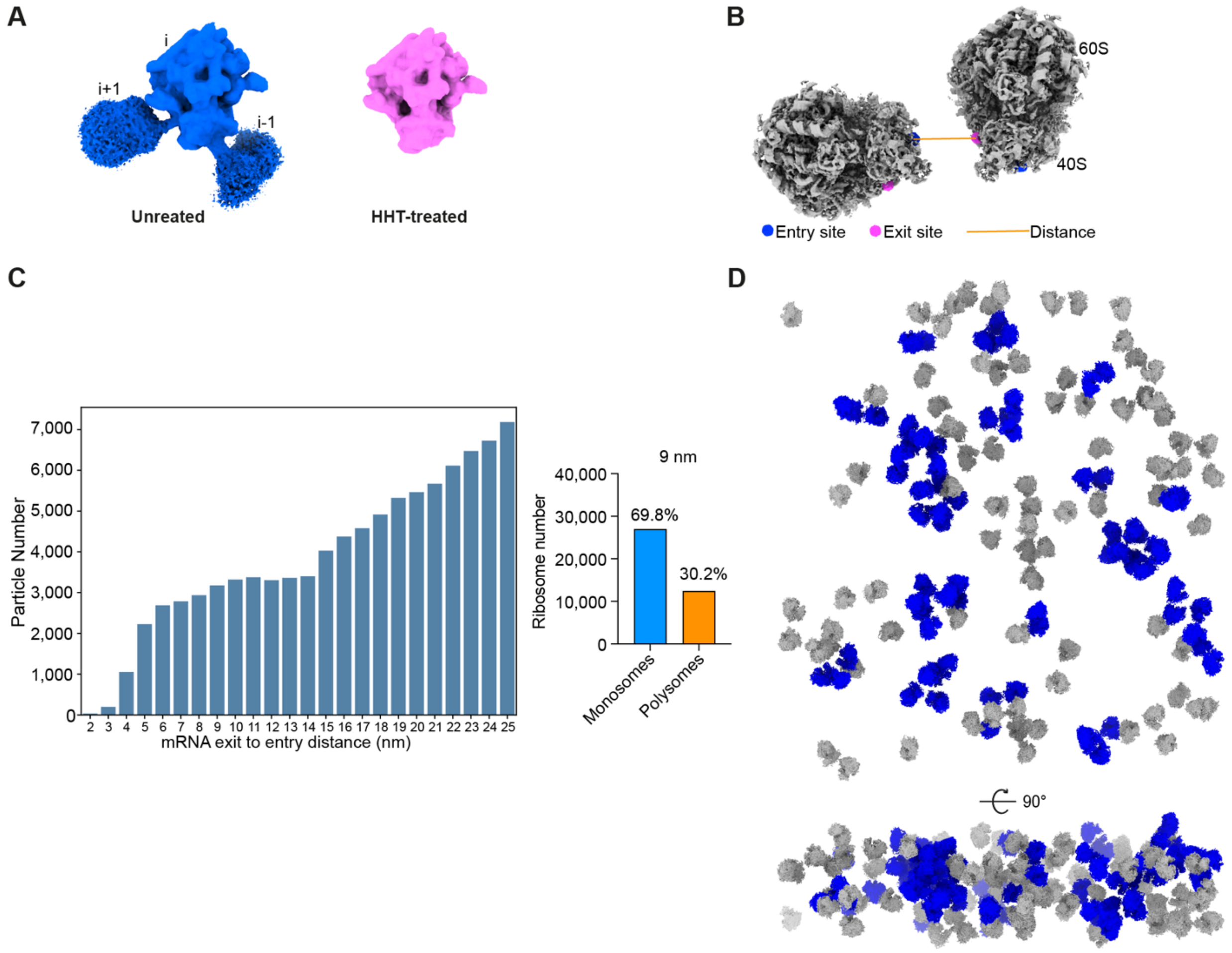


**Fig. S12. Polysome analysis in the untreated dataset.**

(**A**) Subtomogram average map of 39,402 untreated ribosomes (left) and 39,070 HHT-treated ribosomes (right) depicted at similar contour level. The treated ribosome can have a little neighboring density at lower contour level. i, the potential leading ribosome. i+1, the potential trailing ribosome. (**B**) Graphic of the polysomes detection method based on the distance from the mRNA exit site of one ribosome to the entry site of the other ribosome (short for exit-to-entry distance). (**C**) The numbers of detected polysomes within 2 to 25 nm (exit-to-entry distance) proximity in untreated cells (left). The abundance of monosomes and polysomes using the cut-off of 9 nm (right). (**D**) A tomogram showing the detected monosomes (grey) and polysomes (blue).

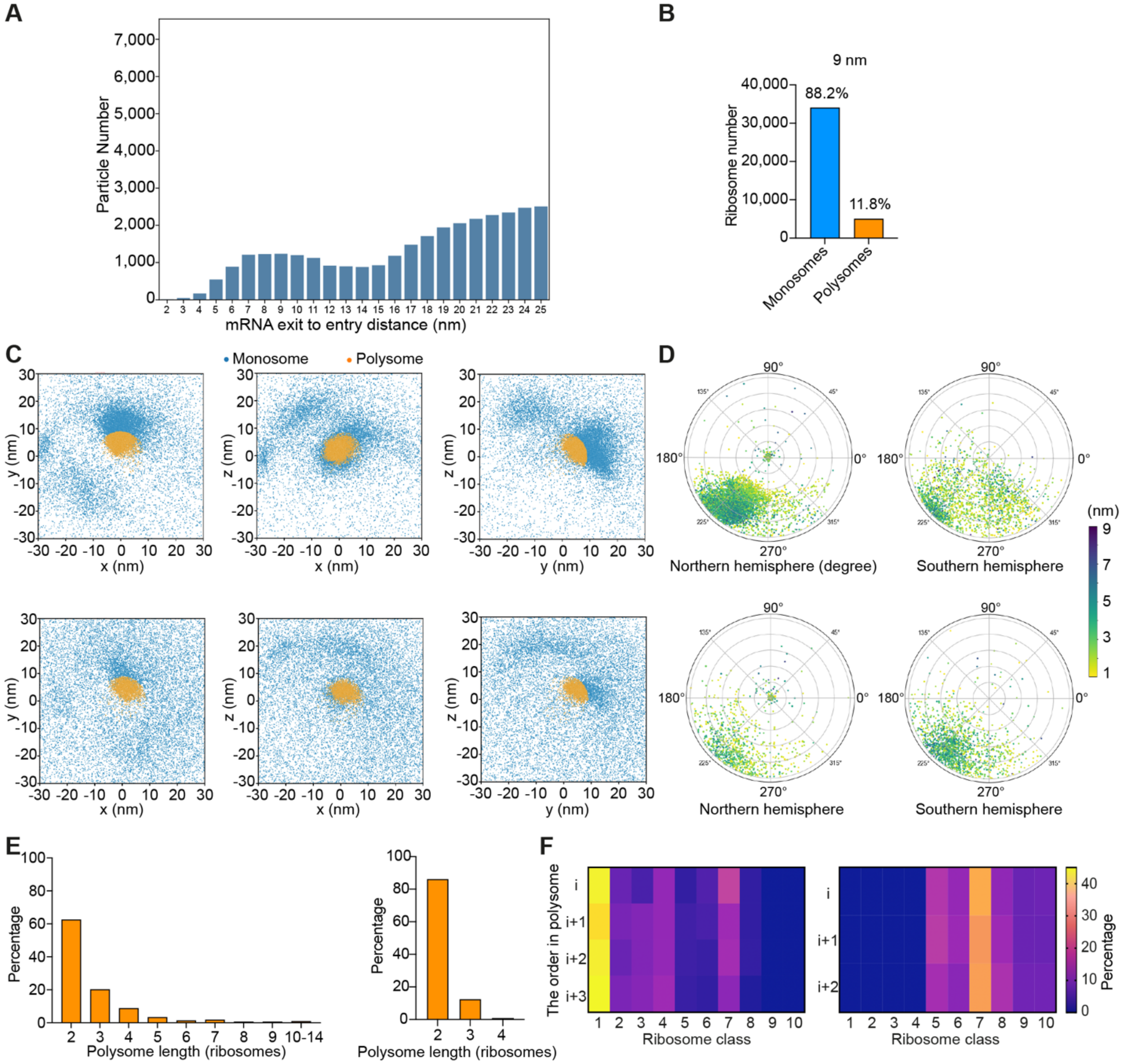


**Fig. S13. Comparison of polysomes between treated and untreated cells.**

(**A**) The numbers of detected polysomes within 2 to 25 nm (exit-to-entry distance) proximity in HHT-treated cells (left). (**B**) The abundance of monosomes and polysomes using the cut-off of 9 nm. (**C**) Distribution of the nearest neighboring ribosomes within 30 nm (exit-to-entry distance) in untreated (top) and treated cells (bottom). The positions of the entry sites of trailing ribosomes were normalized to the exit sites of the leading ribosome positions (these correspond to 0 in all dimensions) and rotated by the inverse rotation of the respective leading ribosomes. The positions corresponding to the ribosomes within polysomes are colored orange, and blue corresponds to monosomes. (**D**) Angular distribution of the trailing ribosomes in polysomes in untreated (top) and treated cells (bottom). The points represent cone rotation (described by Euler angles $\theta$ and $\psi$) of vector (0, 0, 1), projected on the northern hemisphere (for rotated vectors with z coordinate > 0) and southern hemisphere (for rotated vectors with z coordinate <= 0) using stereographic projection. The north pole corresponds to zero rotation, i.e. to a vector (0, 0, 1). The rotations of trailing ribosomes were multiplied by the inverse rotations of the respective leading ribosomes. The points are color-coded based on the Euclidean distance between the leading ribosome exit site and the trailing ribosome entry site. (**E**) Distribution of polysome length in untreated (left) and treated cells (right). (**F**) The ribosome states distribution in i, i+1, i+2 or i+3 in polysomes. i, the leading ribosome. i+n, the trailing ribosome. Left, untreated cells. Right, treated cells. The ribosome class is the same as in Fig. 3A.

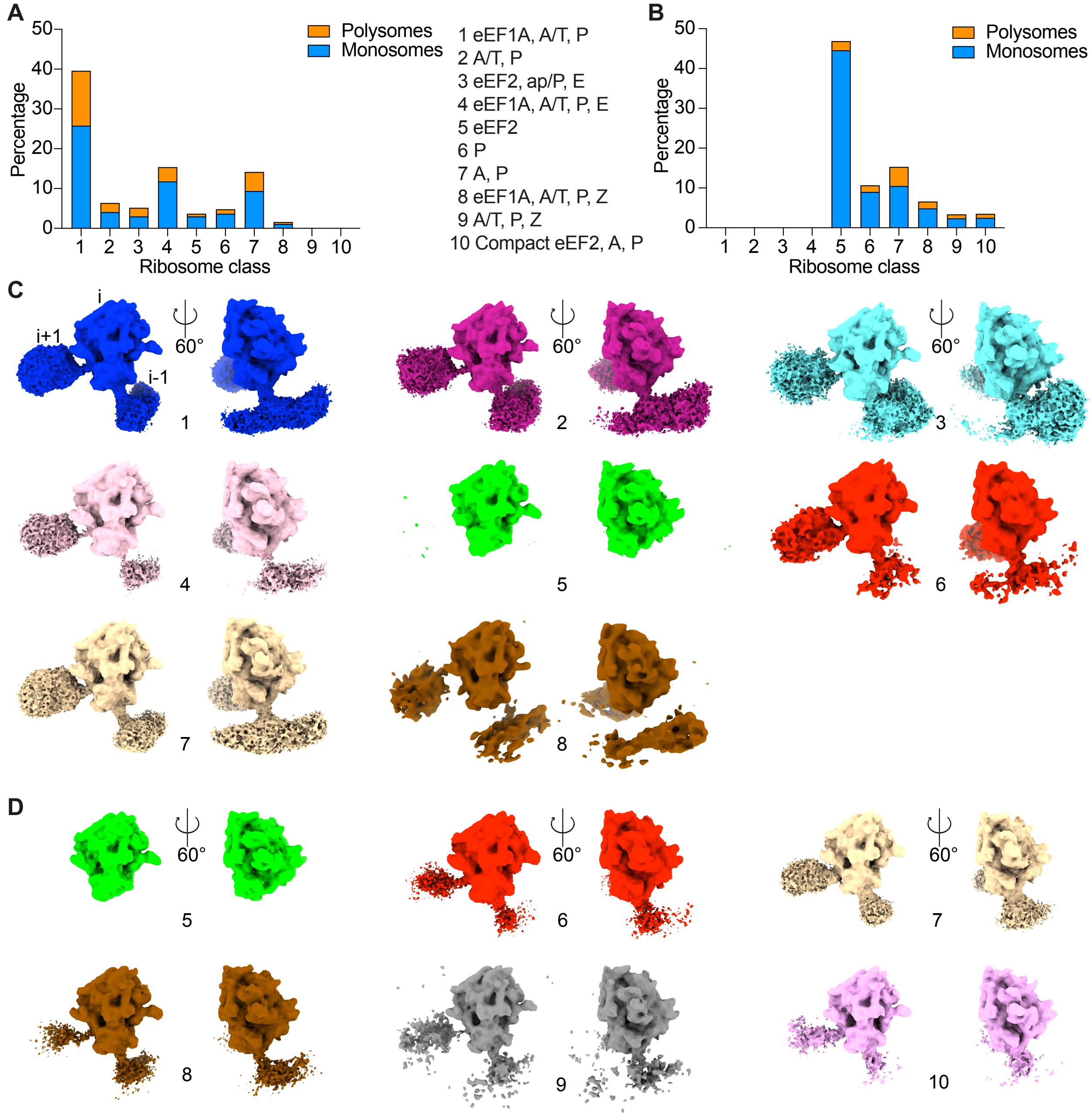


**Fig. S14. The neighborhood density of the individual ribosome state.**

(**A** and **B**) The abundance of mono-ribosomes and polysomes in each state in untreated cells (A) and HHT-treated cells (B). (**C** and **D**) Gaussian-filtered (sDev = 4) filtered ribosome maps of all detected classes in untreated cells (C) and treated cells (D). The class numbers are shown at the bottom and are same as in (A). i, the leading ribosome. i+1, the potential trailing ribosome.

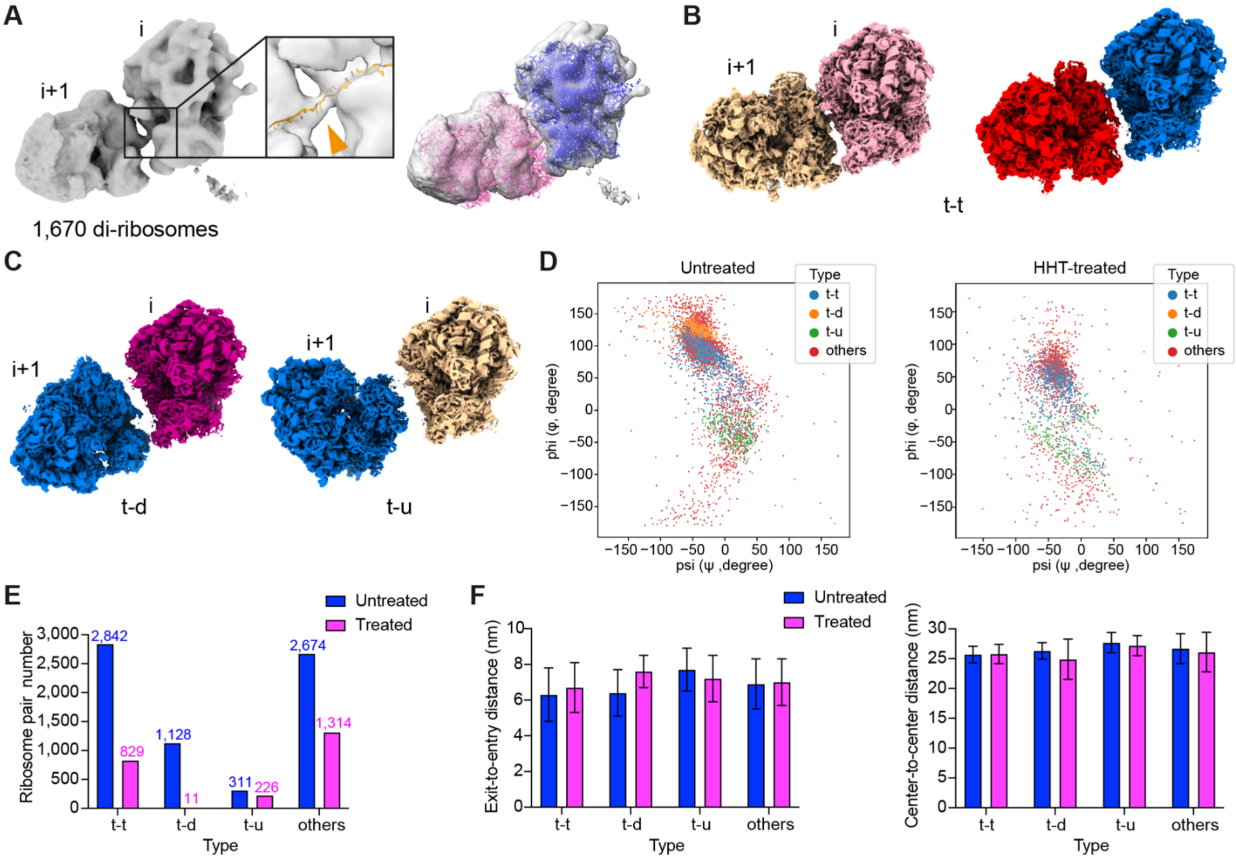


**Fig. S15. Analysis of ribosome pairs in polysomes.**

(**A**) The subtomogram average map of di-ribosome in untreated cells. The disome model (PDB 6I7O) is rigid-body-fitted into the map. For clarity, only mRNA model is shown in orange (left panel). Triangle, mRNA density. i+1, the trailing ribosome. Human collided disome model (PDB 7QVP) is fitted into the map (right panel). Blue and pink represent the two ribosome structures in the disome model. (**B**) Two representative 't-t' ribosome pairs. The translation states of the ribosome pairs are color coded as in Fig. 4. (**C**) Different preferred assembly of adjacent ribosomes in polysomes. t-d, the central protuberance of the 'i+1' ribosome towards down. t-u, the central protuberance of the 'i+1' ribosome towards up. (**D**) The distribution of t-t, t-d and t-u neighboring pairs within polysomes in untreated (left) and treated cells (right). Ψ and φ of trailing ribosomes' orientations were normalized to the zero rotations of the respective leading ribosomes (Materials and methods). (**E**) The abundance of different configurations of ribosome pairs in polysomes. (**F**)

Exit-to-entry distance (left) and center-to-center distance (right) of different types of ribosome pairs in untreated and treated cells. The data represent the mean ± SD. Particle numbers are shown in (E).

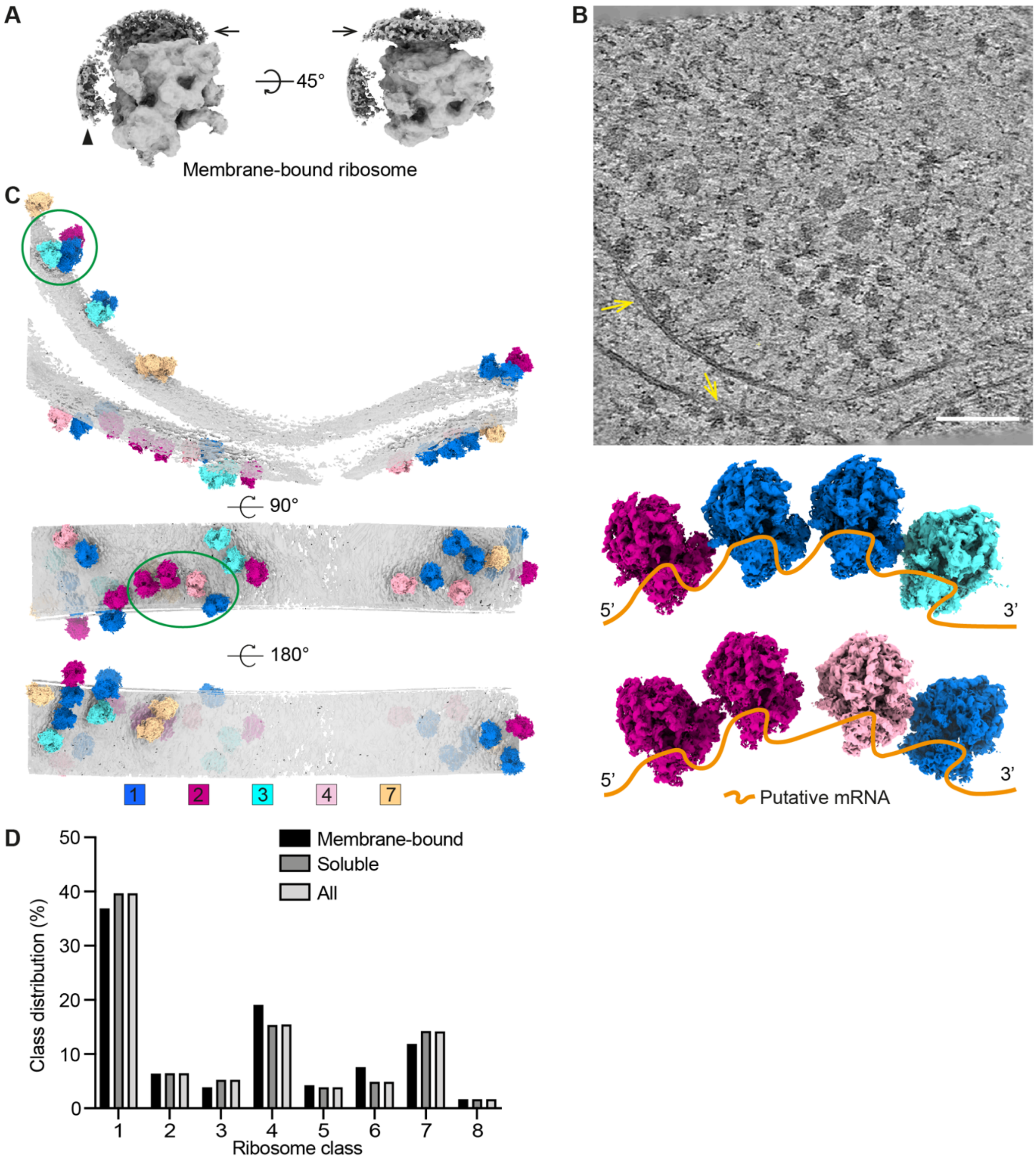


**Fig. S16. Membrane-bound ribosomes inside untreated human cells.**

(**A**) The sub-tomogram average map of membrane-bound ribosomes in untreated cells. The black triangle shows the potential neighboring ribosome. Black arrow, membrane. (**B**) A representative tomographic slice shows some ribosomes (yellow arrow) binding on the membrane. Scale bar, 100nm. (**C**) A segmented membrane with ribosomes from the tomogram in (B). The segmented membrane and the translation states are color coded as indicated in the color bar. Two potential membrane-bound polysomes are circled in green (left panel) and reveal the spatial organization and different translation states (right panel). Orange line, putative mRNA. (**D**) The state distribution of membrane-associated and soluble ribosomes (Materials and methods). The classes (1 to 8) are the same as in Fig. 3.

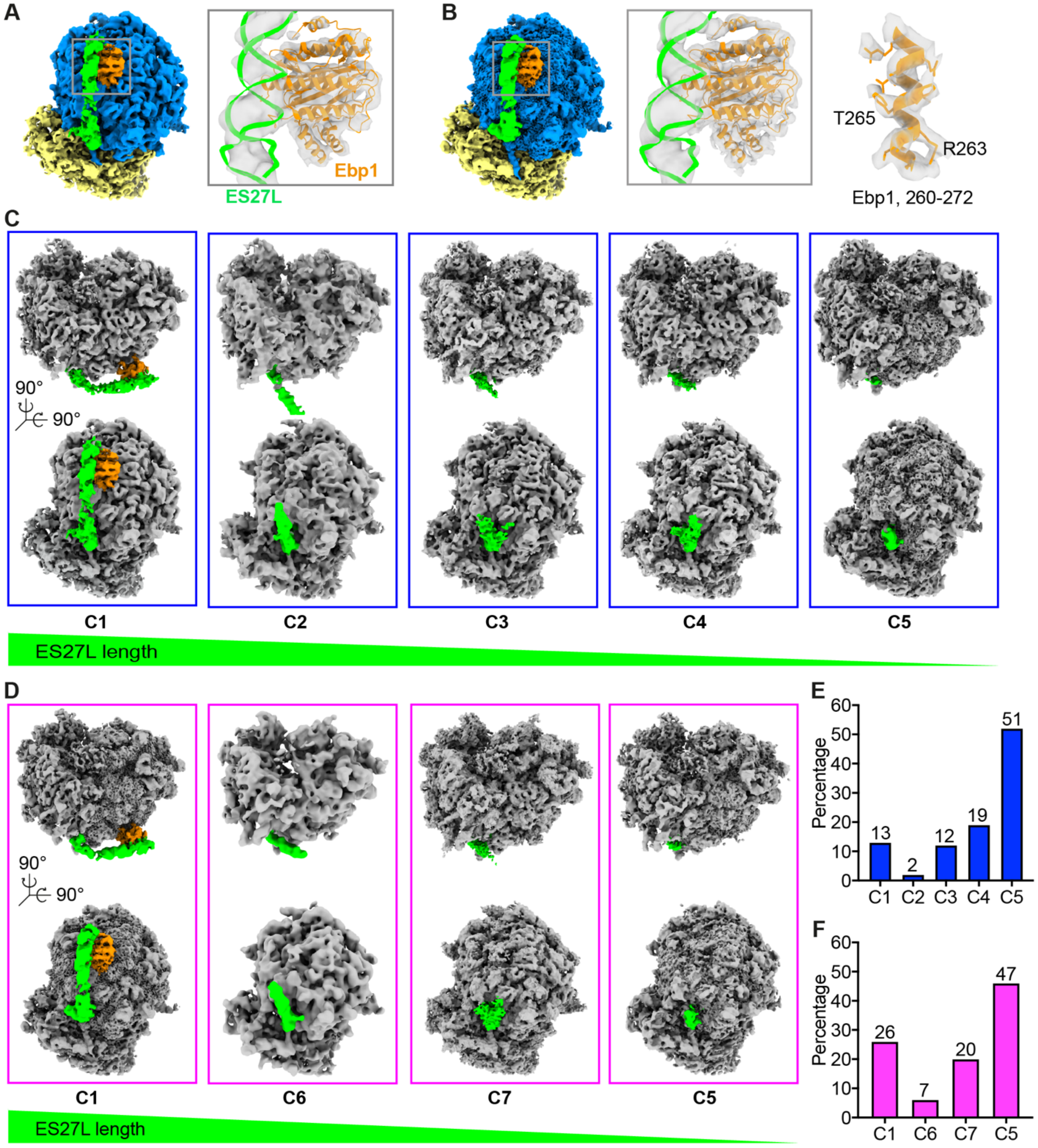


**Fig. S17. Structures of ribosome-associated with Ebp1 and ES27L.**

(**A** and **B**) Structures of ribosomes bound with Ebp1 from untreated cells (A) and HHT-treated cells (B). PDB 6SXO is fitted into the above maps. (**C**) Different conformations of expansion segment ES27L from the untreated cells. C1 to C5: ribosome conformation 1 to 5. The order of

C1 to C5 is defined by the ES27L length. (**D**) Conformations of expansion segment ES27L from the treated dataset. (**E** and **F**) Abundance of C1 to C7 from the two datasets (see tables S2 and S3). Blue, percentage of 39,402 untreated ribosomes. Magenta, percentage of 39,070 treated ribosomes.

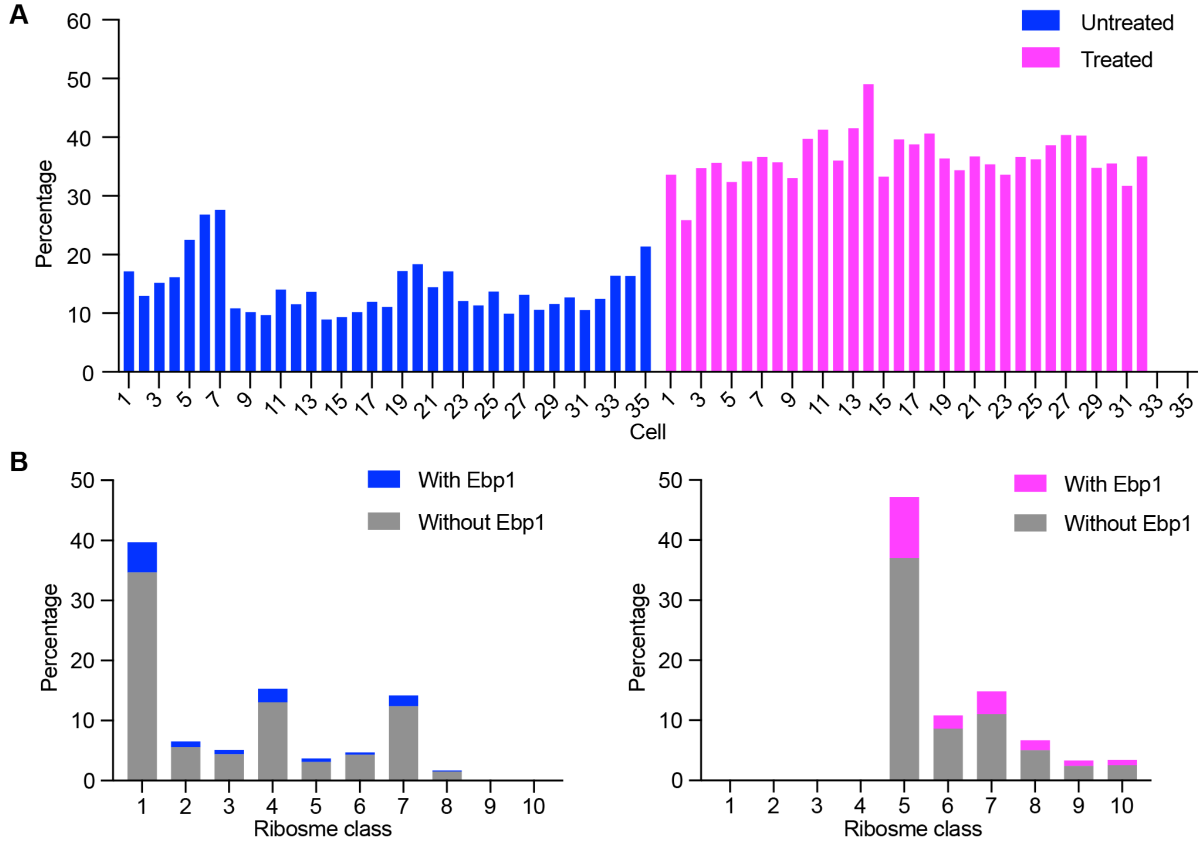


**Fig. S18. Translation states and cellular distribution of Ebp1-associated ribosomes.**

(**A**) Percentage of ribosomes decorated with Ebp1 in 35 untreated and 32 HHT-treated cells. (**B**) Distribution of ribosomes with or without Ebp1 in each ribosome state (the class number is the same as Fig. 3A or fig. S14A). Right, untreated cells. Left, treated cells.

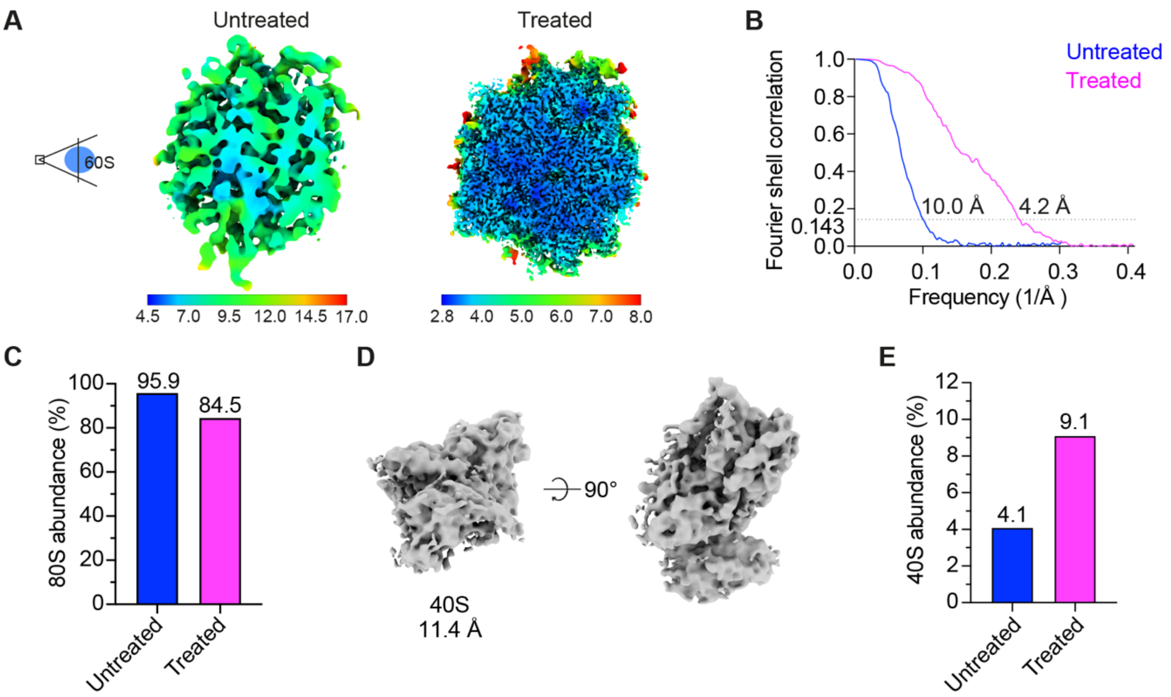


**Fig. S19. Structures of 60S and 40S inside human cells.**

(**A**) Untreated 60S (left) and treated 60S (right) are colored by local resolution calculated in M. Color keys are shown in the bottom. (**B**) FSC curves of 60S from the two datasets (FSC = 0.143). (**C**) Normalized percentage of the 80S ribosome. Abundance = 80S/(60S+80S). (**D**) Subtomogram average map of free 40S in the cytoplasm of untreated cells. (**E**) Percentage of the 40S in untreated and HHT-treated cells, normalized to the number of 80S ribosomes in the respective dataset. Abundance = 40S/(40S+80S).

**Table S1. Cryo-EM data collection of the untreated and treated datasets.**

| | Untreated | HHT-treated |
|---|---|---|
| Microscope | Titan Krios G4 | Titan Krios G4 |
| Voltage (kV) | 300 | 300 |
| Camera | Falcon4 | Falcon4 |
| Magnification | 105,000 | 105,000 |
| Pixel size (Å) | 1.223 | 1.223 |
| Defocus range (µm) | -1.5 to -4.5 | -1.5 to -4.5 |
| Automation software | SerialEM | SerialEM |
| Energy filter slit width (eV) | 10 | 10 |
| Electron exposure (e/Å$^2$) | 120 to 150 | 120 to 150 |
| Number of tilt series | 358 | 352 |

**Table S2. Cryo-EM data refinement and validation statistics in untreated cells.**

| | Symmetry imposed | Initial particle number | Final particle number | Resolution (Å) FSC =0.143 | Resolution range(Å) | Global B factor ($Å^2$) |
|---|---|---|---|---|---|---|
| 80S ribosome<br>EMD-16721 | C1 | 286,400 | 39,402 | 3.2 | 2.5 to 7.5 | -65 |
| 80S-eEF1A, A/T, P<br>EMD-16725 | C1 | 286,400 | 15,587 | 3.4 | 2.7 to 8 | -48 |
| 80S-A/T, P<br>EMD-16726 | C1 | 286,400 | 2,538 | 8.3 | 4.0 to 12 | -74 |
| 80S-eEF2, ap/P, E<br>EMD-16727 | C1 | 286,400 | 2,037 | 9.6 | 5.1 to 13 | -153 |
| 80S-eEF1A, A/T, P, E<br>EMD-16728 | C1 | 286,400 | 6,054 | 8.7 | 4.1 to 12 | -133 |
| 80S-eEF2<br>EMD-16733 | C1 | 286,400 | 1,495 | 11.7 | 7.7 to 15 | -200 |
| 80S-P<br>EMD-16734 | C1 | 286,400 | 1,892 | 14.7 | 10.6 to 20 | -200 |
| 80S-A, P<br>EMD-16735 | C1 | 286,400 | 5,574 | 5.0 | 3.1 to 8 | -77 |
| 80S-eEF1A, A/T, P, Z<br>EMD-16736 | C1 | 286,400 | 636 | 16.4 | 9.7 to 20 | -200 |
| Di-ribosome<br>EMD-16737 | C1 | 286,400 | 1,607 | 18.1 | 12 to 30 | -200 |
| 80S<br>ES27L conformation 1<br>EMD-16738 | C1 | 286,400 | 5,142 | 6.6 | 3.2 to 15 | -60 |
| 80S<br>ES27L conformation 2<br>EMD-16739 | C1 | 286,400 | 832 | 13.7 | 10 to 20 | -200 |
| 80S<br>ES27L conformation 3<br>EMD-16740 | C1 | 286,400 | 4,715 | 6.9 | 3.5 to 15 | -78 |
| 80S<br>ES27L conformation 4<br>EMD-16741 | C1 | 286,400 | 7,332 | 4.4 | 2.9 to 13 | -49 |
| 80S<br>ES27L conformation 5<br>EMD-16742 | C1 | 286,400 | 19,902 | 3.6 | 2.8 to 10 | -59 |
| Free 60S<br>EMD-16743 | C1 | 286,400 | 1,693 | 10.0 | 4.5 to 17 | -178 |

**Table S3. Cryo-EM data refinement and validation statistics in HHT-treated cells.**

| | Symmetry imposed | Initial particle number | Final particle number | Resolution (Å) FSC =0.143 | Resolution range(Å) | Map sharpening B factor (Å$^2$) |
|---|---|---|---|---|---|---|
| 80S ribosome<br>EMD-16722 | C1 | 281,600 | 39,070 | 3.2 | 2.5 to 7.5 | -46 |
| 80S-eEF2<br>EMD-16744 | C1 | 281,600 | 18,320 | 3.7 | 2.7 to 8 | -66 |
| 80S-P<br>EMD-16748 | C1 | 281,600 | 4,198 | 8.5 | 3.7 to 12 | -85 |
| 80S-A, P<br>EMD-16747 | C1 | 281,600 | 5,963 | 4.4 | 2.8 to 8 | -58 |
| 80S-eEF1A, A/T, P, Z<br>EMD-16749 | C1 | 281,600 | 2,607 | 8.2 | 3.7 to 12 | -91 |
| 80S-A/T, P, Z<br>EMD-16750 | C1 | 281,600 | 1,301 | 11.5 | 6.7 to 14 | -200 |
| 80S-Compact eEF2, A, P<br>EMD-16751 | C1 | 281,600 | 1,344 | 11.1 | 6.3 to 14 | -200 |
| 80S<br>ES27L conformation 1<br>EMD-16752 | C1 | 281,600 | 10,249 | 4.0 | 2.8 to 13 | -59 |
| 80S<br>ES27L conformation 5<br>EMD-16754 | C1 | 281,600 | 18,344 | 3.8 | 2.8 to 12 | -63 |
| 80S<br>ES27L conformation 6<br>EMD-16755 | C1 | 281,600 | 2,625 | 11.0 | 5.8 to 20 | -245 |
| 80S<br>ES27L conformation 7<br>EMD-16756 | C1 | 281,600 | 7,852 | 4.1 | 2.9 to 12 | -56 |
| Free 60S<br>EMD-16757 | C1 | 281,600 | 7,176 | 4.2 | 2.8 to 8 | -45 |

**Table S4. Pseudocode for polysome chain tracing.**

**Algorithm: Trace polysome chains in a single tomogram**

```
untraced_ribosomes = ribosomes
traced_ribosomes = empty list
for each ribosome do
        if ribosome is in untraced_ribosomes then
                trace_chain = true
                current_chain = ribosome
                remove ribosome from untraced_ribosomes
                while trace_chain is true do
                        NN = nearest neighbor in untraced_ribosomes
                        if the distance to the NN is within the range then
                                current_chain = add NN
                                remove NN from untraced_ribosomes
                        else
                                trace_chain = false
                                for the first ribosome in current_chain do:
                                        NN = nearest neighbor in traced_ribosomes
                                        if the distance to the NN is within the range:
                                                append current_chain to the existing one
                                for the last ribosome in current_chain do:
                                        NN = nearest neighbor in traced_ribosomes
                                        if the distance to the NN is within the range:
                                                prepend current_chain to the existing one
                                traced_ribosomes = add current_chain ribosomes
                        end if
                end while
        end if
end for
```

**Movie S1.**

A tomogram and the template matching of 80S ribosomes.  For visualization, the tomogram was set to 50% transparency, and the CCC value of 80S was colored in yellow.

**Movie S2.**

Three-dimensional view of HHT and the neighboring 28S rRNA from the HHT-treated human cells.

**Movie S3.**

Close view of the Z t-RNA at the ‘eEF1A, A/T, P, Z’ state in HHT-treated cells. Tangerine, Z-tRNA (PDB 6MTB).
